# Whole transcriptome sequencing reveals drought resistance-related genes in upland cotton

**DOI:** 10.1101/2021.11.11.468302

**Authors:** Juyun Zheng, Zeliang Zhang, Yajun Liang, Zhaolong Gong, Zhiwei Sang, Xueyuan Li, Jungduo Wang

**Author notes:** These authors have contributed equally to this work (Juyun Zheng, Zeliang Zhang, Yajun Liang).

## Abstract

China, especially the Xinjiang cotton area, is facing severe agricultural water shortages, which seriously restrain the development of the cotton industry. Discovering cotton drought resistance genes and cultivating high-quality and drought-resistant cotton materials through molecular breeding methods are of great significance to the development of the cotton industry. In this study, the drought-resistant cotton material Xinluzhong NO.82 and the drought-sensitive cotton material Kexin NO.1 were used to identify a batch of drought-resistant candidate genes through whole transcriptome sequencing. The main research results obtained were as follows: the ceRNA (competing endogenous RNAs) network was constructed using full transcriptional sequencing to screen the core genes in the core pathway; two drought-related candidate genes were obtained. Gohir.A11G156000 was upregulated at 0 h vs 12 h and downregulated at 12 h vs 24 h. Gohir.A07G220600 was downregulated at 0 h vs 12 h and upregulated at 12 h vs 24 h. The results for drought-resistant materials and drought-sensitive materials were similar. Gohir.A11G156000, encoding GABA-T, which is homologous to POP2 in Arabidopsis thaliana, affects the drought resistance of plants by regulating the GABA content. Gohir.A07G220600 encodes L-aspartate oxidase, which is homologous to AO in Arabidopsis thaliana, and is involved in the early steps of NAD biosynthesis and in plant antioxidant reactions. This study confirmed that the use of gene expression regulatory networks can quickly screen reliable drought-resistance genes and can be used for subsequent gene function verification.

## 1. Introduction

Drought is a worldwide problem that contributes to the abiotic stress of plants throughout their lifecycle; the effects of water deficiency on crops are complex and variable, which can delay plant development and thus affect plant morphology as well as physiology (Mehta et al.,2017; Do et al.,2013). Plants inhabiting drought-prone areas have, however, developed various strategies to cope with stress, including developing larger and deeper root systems to increase water absorption from the deep soil, regulating stomatal closure to reduce water loss, accumulating compatible solutes and protective proteins and increasing the level of antioxidants (Chaves et al.,2003). With the continuous development of reverse resistance genetic engineering technology, transgenic technology can be used to introduce reverse resistance genes into plants to obtain new materials. This not only alleviates the problem that cotton competes with food crops for water resources but also increases farmers’ income, guarantees cotton yield, and promotes continuous development of the textile industry.

In recent years, there has been an increasing number of studies on plant drought resistance mechanisms using transcriptome sequencing, and a large number of genes related to drought resistance have been mined (Raney et al.,2014;Bowman et al.,2013;Thumma et al.,2012). Transcriptome sequencing makes it possible to obtain, as a whole, transcripts for the expression of all genes in a certain organism under specific physiological conditions or treatments so that the functions and structures of genes can be studied at a global level. Transcriptome sequencing has become a powerful tool for studying plant drought resistance mechanisms and mining genes underlying drought resistance (Luo et al.,2009;Deyholos et al.,2010). Bowman et al. (2013) used RNA-Seq to profile natural rain-fed and well-watered cotton in the field, and a total of 1530 differential transcripts were coexpressed in well-irrigated and drought-stressed root tissues. Chen et al. (2013), through transcriptome sequencing, found that upregulated genes in cotton under drought stress were mainly involved in glycerophospholipid metabolism, glycolysis/gluconeogenesis, amino and nucleotide sugar metabolism, lysosome, alanine, aspartate and glutamate metabolism, fatty acid metabolism, pyruvate metabolism, and galactose metabolism pathways, such as cysteine and methionine metabolism. The downregulated genes were mainly involved in the photosynthesis, photosynthetic antenna proteins, glyceride metabolism, oxidative phosphorylation, glycolysis/gluconeogenesis, phytohormone signalling, flavonoid biosynthesis, porphyrin and chlorophyll metabolism, and nitrogen metabolism pathways.

Zhang et al. (2015), through transcriptome sequencing, showed that moderate drought of liquorice could inhibit the expression of multiple enzymes in the cell wall and promote the synthesis pathways of terpenoids and flavonoids, which in turn revealed the molecular mechanism of liquorice adaptation to the accumulation of active components during drought. Liu et al. (2015) found that upregulated genes in rape roots under drought stress were mainly involved in stimulated and stress-related biological processes by transcriptome sequencing, while upregulated genes in leaves mainly played the roles in cells and cellular components. In addition, the regulatory network of transcription factors under drought stress was analysed. Chen et al. (2016) investigated the transcripts of soybean under drought conditions using RNA-Seq, and the results showed that drought inhibited photosynthesis and chlorophyll synthesis and promoted cell wall synthesis. Rohini Garg et al. (2016) found that the differentially expressed genes in chickpeas under drought stress were mainly involved in photosynthesis, trehalose synthesis, citrulline synthesis, UDP glucose synthesis and other pathways.

In this study, we used laboratory-reserved varieties Xinluzhong No. 82 and Kexin No. 1 to obtain three-leaf cotton seedlings by culturing for 28 days through hydroponic experiments. The plants were subjected to drought stress experiments with 17% PEG (polyethylene glycol) solution. At 0 h, 12 h, and 24 h, the leaves were taken for full transcriptome analysis to obtain differentially expressed genes of different varieties and processed at different time points, and the accuracy of the whole transcriptome results was verified by qRT-PCR (real-time fluorescent quantitative PCR). GO clustering and KEGG pathway analysis of differentially expressed genes were performed, ceRNA (competing endogenous RNAs) pathways were constructed and analysed, existing literature reports were synthesized, cotton drought-resistant pathways were screened, and key genes in key pathways were obtained. Homologous sequence comparison with Arabidopsis sequences was performed to finally obtain candidate drought resistance genes, which provided a theoretical basis for the isolation and identification of stress resistance-related genes in cotton and other crops.

## 2. Results

### 2.1 Total RNA quality control results

After sequencing quality control, a total of 307.23 Gb of clean read data was obtained. The Q30 base percentage of each sample was not less than 93.73%, and the GC content was not less than 42.0%. The comparison efficiency between the reads of each sample and the reference genome ranged from 95.67% to 96.92% (Table 1), which suggests that the data can be used for subsequent analysis.

**Table 1.** Statistical table of sequencing data evaluation

| SampleID | ReadSum | BaseSum | GC(%) | N(%) | Q20(%) | Q30(%) | Total Reads | Mapped Reads |
| --- | --- | --- | --- | --- | --- | --- | --- | --- |
| XJ0hR1 | 71742474 | 21224569062 | 42.85 | 0 | 97.92 | 93.80 | 143484948 | 138314357(96.40%) |
| XJ0hR2 | 53952826 | 16058270844 | 42.78 | 0 | 97.93 | 93.88 | 107905652 | 104104314(96.48%) |
| XJ0hR3 | 67100443 | 19944130312 | 43.07 | 0 | 97.90 | 93.73 | 134200886 | 129284996(96.34%) |
| XJ0hS1 | 53931117 | 16041131562 | 42.77 | 0 | 98.34 | 94.70 | 107862234 | 103981706(96.40%) |
| XJ0hS2 | 57233313 | 17014361012 | 42.72 | 0 | 98.32 | 94.65 | 114466626 | 110557335(96.58%) |
| XJ0hS3 | 56289536 | 16740674196 | 42.84 | 0 | 98.21 | 94.50 | 112579072 | 108590412(96.46%) |
| XJ12hR1 | 60305370 | 17970064928 | 43.00 | 0 | 98.27 | 94.52 | 120610740 | 116216676(96.36%) |
| XJ12hR2 | 53171614 | 15800217442 | 42.67 | 0 | 98.28 | 94.59 | 106343228 | 102417327(96.31%) |
| XJ12hR3 | 54393796 | 16198260692 | 42.06 | 0 | 98.29 | 94.60 | 108787592 | 104988226(96.51%) |
| XJ12hS1 | 55632640 | 16553339614 | 42.5 | 0 | 98.27 | 94.51 | 111265280 | 107405318(96.53%) |
| XJ12hS2 | 55334885 | 16503777878 | 42.61 | 0 | 98.19 | 94.39 | 110669770 | 105879284(95.67%) |
| XJ12hS3 | 56241086 | 16733164858 | 42.56 | 0 | 98.20 | 94.39 | 112482172 | 107971513(95.99%) |
| XJ24hR1 | 55174614 | 16453704450 | 42.76 | 0 | 98.21 | 94.48 | 110349228 | 106142505(96.19%) |
| XJ24hR2 | 55843015 | 16660958134 | 42.43 | 0 | 98.34 | 94.76 | 111686030 | 107264860(96.04%) |
| XJ24hR3 | 58592920 | 17464204712 | 42.98 | 0 | 98.22 | 94.44 | 117185840 | 113571833(96.92%) |
| XJ24hS1 | 55363695 | 16530287984 | 42.26 | 0 | 98.47 | 94.98 | 110727390 | 106999856(96.63%) |
| XJ24hS2 | 55231921 | 16459828202 | 42.49 | 0 | 98.19 | 94.32 | 110463842 | 106313930(96.24%) |
| XJ24hS3 | 56920057 | 16913326528 | 42.8 | 0 | 98.45 | 94.91 | 113840114 | 109889372(96.53%) |

Note: SampleID: sample name. ReadSum: the total number of paired-end reads in the clean data. BaseSum: Total base number of the clean data. GC (%): GC content, namely, the percentage of G and C bases in the total
bases in the clean data. N (%): Unresolved base percentage of total bases in the clean data. Q30 (%): Percentage of bases whose clean data mass value is greater than or equal to Q30. Total Reads: the number of clean reads, counted as single-ended; Mapped Reads: the number of reads compared to the reference genome and the percentage in clean reads.

### 2.2 Analysis of sequencing results for the cotton transcriptome

The FASTQ data obtained by high-throughput sequencing were used to predict and identify the four RNA types. The total number of RNAs was counted and included known RNAs such as mRNA81336, lncRNA6906, miRNA453, and circRNA948. By dimension reduction of the principal component analysis, the distribution of the sample points was shown on a two- or three-dimensional plane. EIGENSOFT 7.2.1 (Alkes et al.2006) software was used to perform principal component analysis to cluster the samples. Samples treated at different times were clearly separated, and the distance between biological duplicates showed better consistency between repetitions and differences between different groups. The first three components explained 89.1%, 4.8%, and 1.9% of the variation. Spearman’s correlation coefficient r (Spearman’s correlation coefficient) was used to evaluate the biological replicates. The closer r2 is to 1, the stronger the correlation between the two repeated samples. Generally, when r2 is greater than 0.9, the correlation between two repeated samples is considered to be good. The results show that the correlation between repeated samples of experimental materials was strong and that the difference between the two materials was large, but the correlations between repeated samples were better (Figure 1).

**Figure 1.**
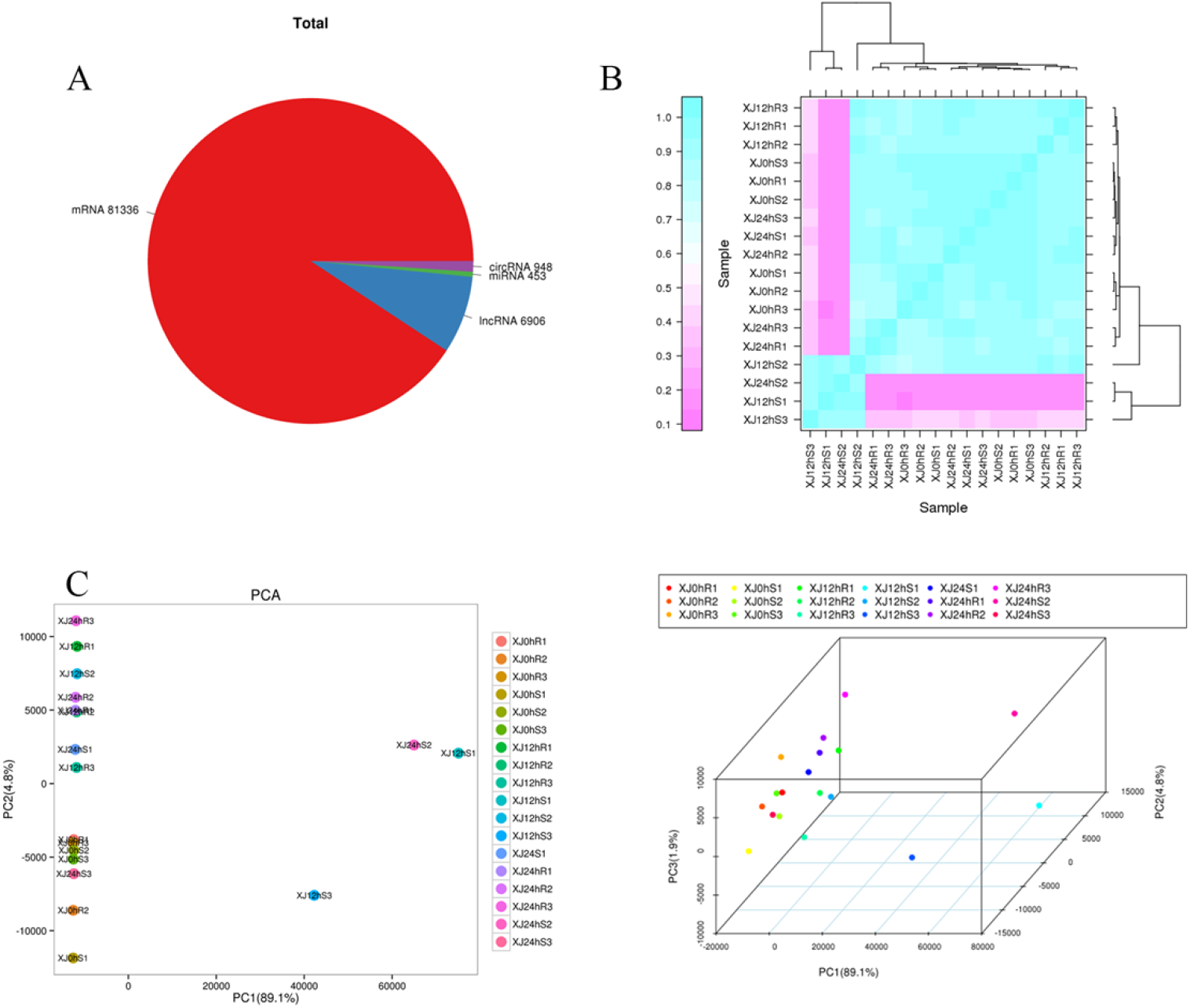
A:Statistics of different RNA quantities, B:a sample correlation diagram, different colours in the figure represent different correlation coefficient values. The abscissa and ordinate represent different samples.C:a two-dimensional PCA clustering map, and a three-dimensional PCA clustering map, the samples are gathered into three dimensions by PCA: PC1 represents the first principal component, PC2 represents the second principal component, and PC3 represents the third principal component, a point represents a sample, and a colour represents a grouping.

### 2.3 Differential gene expression analysis

#### 2.3.1 Differential gene expression screening

Based on the sequencing results, many differentially expressed genes were obtained. Using |log2(fold change)|>1, fold change ≥ 1.5 and P value <0.01 as the screening criteria, significant differentially expressed genes were identified. According to the gene expression levels in different samples, 26,923 differentially expressed genes were identified. Most of the comparisons revealed more downregulated genes than upregulated genes. The differentially expressed genes from the same material processed at different times were far greater than the differentially expressed genes from different materials processed at the same time, and the statistics of the number of differentially expressed genes are as follows (Table 2):

**Table 2.**
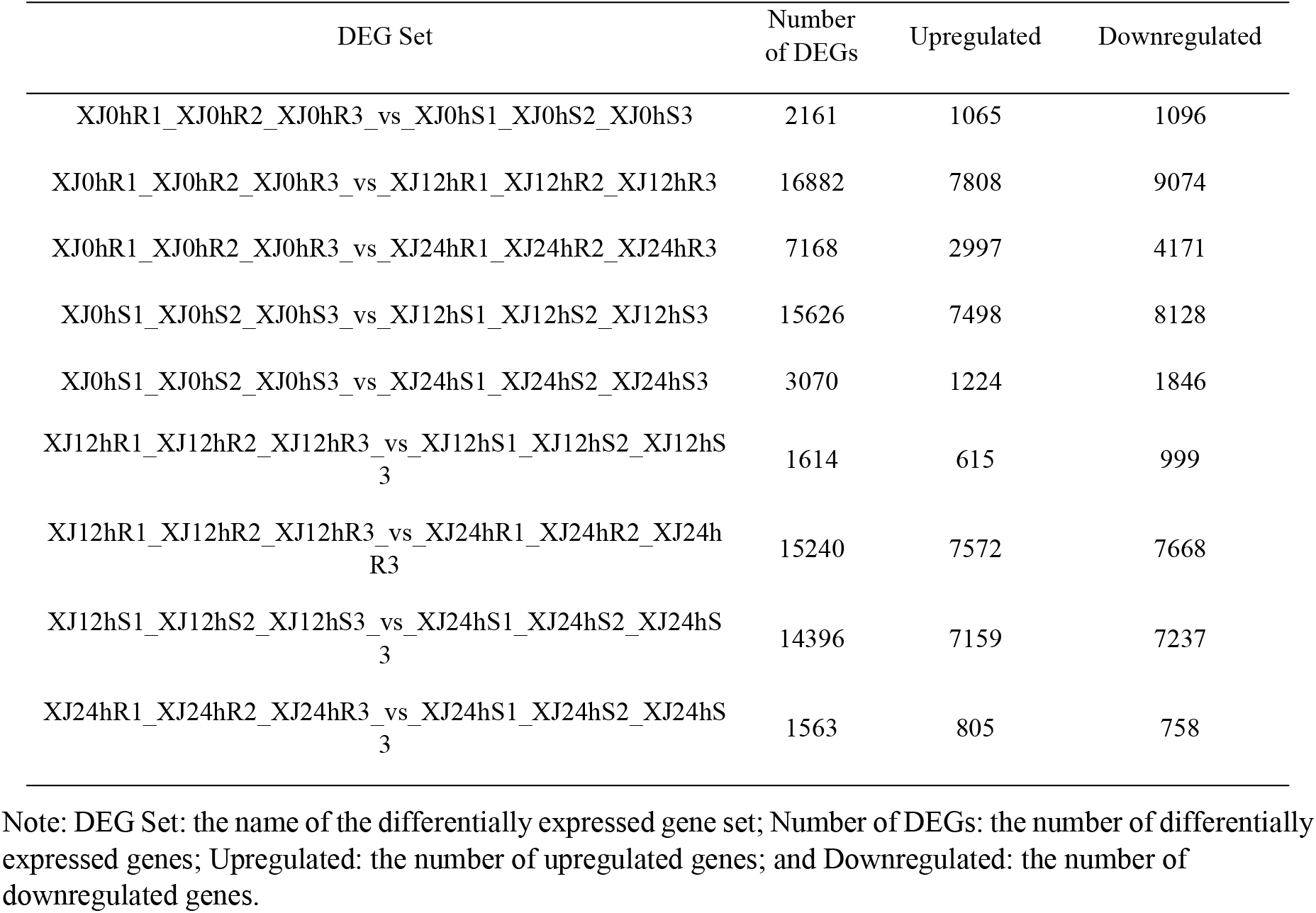
Statistics of the number of differentially expressed genes

| DEG Set | Number of DEGs | Upregulated | Downregulated |
| --- | --- | --- | --- |
| XJ0hR1_XJ0hR2_XJ0hR3_vs_XJ0hS1_XJ0hS2_XJ0hS3 | 2161 | 1065 | 1096 |
| XJ0hR1_XJ0hR2_XJ0hR3_vs_XJ12hR1_XJ12hR2_XJ12hR3 | 16882 | 7808 | 9074 |
| XJ0hR1_XJ0hR2_XJ0hR3_vs_XJ24hR1_XJ24hR2_XJ24hR3 | 7168 | 2997 | 4171 |
| XJ0hS1_XJ0hS2_XJ0hS3_vs_XJ12hS1_XJ12hS2_XJ12hS3 | 15626 | 7498 | 8128 |
| XJ0hS1_XJ0hS2_XJ0hS3_vs_XJ24hS1_XJ24hS2_XJ24hS3 | 3070 | 1224 | 1846 |
| XJ12hR1_XJ12hR2_XJ12hR3_vs_XJ12hS1_XJ12hS2_XJ12hS3 | 1614 | 615 | 999 |
| XJ12hR1_XJ12hR2_XJ12hR3_vs_XJ24hR1_XJ24hR2_XJ24hR3 | 15240 | 7572 | 7668 |
| XJ12hS1_XJ12hS2_XJ12hS3_vs_XJ24hS1_XJ24hS2_XJ24hS3 | 14396 | 7159 | 7237 |
| XJ24hR1_XJ24hR2_XJ24hR3_vs_XJ24hS1_XJ24hS2_XJ24hS3 | 1563 | 805 | 758 |

Note: DEG Set: the name of the differentially expressed gene set; Number of DEGs: the number of differentially expressed genes; Upregulated: the number of upregulated genes; and Downregulated: the number of downregulated genes.

#### 2.3.2 Differential lncRNA expression screening

In the process of detecting differentially expressed lncRNAs, fold change represents the ratio of expression levels between two samples (groups). After high-throughput sequencing, 6906 lncRNAs were identified, with 1,639 differentially expressed lncRNAs. The number of differentially expressed lncRNAs in each group was counted as shown in the table below (Table 3).

**Table 3.** Statistics of the number of differentially expressed lncRNAs

| DEG Set | Number of DEGs | Upregulated | Downregulated |
| --- | --- | --- | --- |
| XJ0hR1_XJ0hR2_XJ0hR3_vs_XJ0hS1_XJ0hS2_XJ0hS3 | 294 | 147 | 147 |
| XJ0hR1_XJ0hR2_XJ0hR3_vs_XJ12hR1_XJ12hR2_XJ12hR3 | 462 | 246 | 216 |
| XJ0hR1_XJ0hR2_XJ0hR3_vs_XJ24hR1_XJ24hR2_XJ24hR3 | 182 | 108 | 74 |
| XJ0hS1_XJ0hS2_XJ0hS3_vs_XJ12hS1_XJ12hS2_XJ12hS3 | 439 | 220 | 219 |
| XJ0hS1_XJ0hS2_XJ0hS3_vs_XJ24hS1_XJ24hS2_XJ24hS3 | 126 | 73 | 53 |
| XJ12hR1_XJ12hR2_XJ12hR3_vs_XJ12hS1_XJ12hS2_XJ12hS3 | 302 | 148 | 154 |
| XJ12hR1_XJ12hR2_XJ12hR3_vs_XJ24hR1_XJ24hR2_XJ24hR3 | 437 | 217 | 220 |
| XJ12hS1_XJ12hS2_XJ12hS3_vs_XJ24hS1_XJ24hS2_XJ24hS3 | 423 | 223 | 200 |
| XJ24hR1_XJ24hR2_XJ24hR3_vs_XJ24hS1_XJ24hS2_XJ24hS3 | 342 | 162 | 180 |

Note: DEG Set: the name of the differentially expressed gene set; Number of DEGs: the number of differentially expressed lncRNAs; Upregulated: the number of upregulated lncRNAs; and Downregulated: the number of downregulated lncRNAs.

Hierarchical clustering analysis was performed on the selected differentially expressed lncRNAs, and lncRNAs with the same or similar expression behaviour were clustered. According to the heat map, most lncRNAs may be constitutively expressed under drought stress and have a tolerance effect (Figure 2).

**Figure 2.**
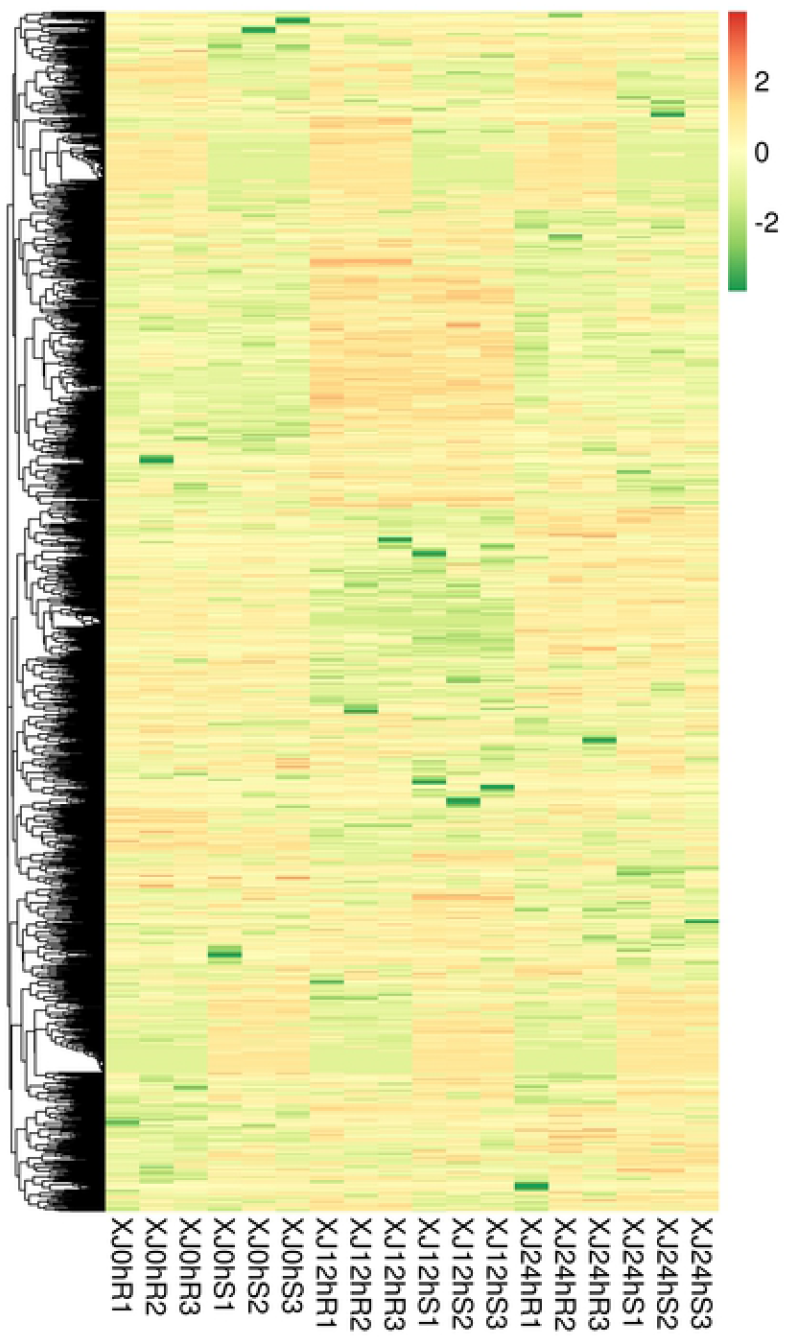
Differentially expressed lncRNA cluster map Note: Different columns in the figure represent different samples, and different rows represent different lncRNAs. The colour represents the expression level of an lncRNA in the sample as log10 (FPKM+0.000001).

#### 2.3.3 Differential circRNA expression screening

In the process of detecting differentially expressed circRNAs, a fold change greater than or equal to 1.5 and a P-value less than 0.05 were used as the screening criteria. Fold change represents the ratio of expression levels between two samples (groups). Since the differential expression analysis of circRNA is an independent statistical hypothesis test on a large number of circRNA expression levels, false positives will be an issue. Therefore, in the analysis process, the p value is used as a key indicator for screening differentially expressed circRNAs. Based on the prediction software, a total of 948 circRNAs were identified, and the number of differential circRNAs was small. The statistics of the number of differentially expressed circRNAs in each group are as follows (Table 4).

**Table 4.** Statistics of the number of differentially expressed circRNAs

| DEG Set | Number of DEGs | Upregulated | Downregulated |
| --- | --- | --- | --- |
| XJ0hR1_XJ0hR2_XJ0hR3_vs_XJ0hS1_XJ0hS2_XJ0hS3 | 8 | 0 | 8 |
| XJ0hR1_XJ0hR2_XJ0hR3_vs_XJ12hR1_XJ12hR2_XJ12hR3 | 11 | 6 | 5 |
| XJ0hR1_XJ0hR2_XJ0hR3_vs_XJ24hR1_XJ24hR2_XJ24hR3 | 1 | 0 | 1 |
| XJ0hS1_XJ0hS2_XJ0hS3_vs_XJ12hS1_XJ12hS2_XJ12hS3 | 2 | 2 | 0 |
| XJ0hS1_XJ0hS2_XJ0hS3_vs_XJ24hS1_XJ24hS2_XJ24hS3 | 6 | 6 | 0 |
| XJ12hR1_XJ12hR2_XJ12hR3_vs_XJ12hS1_XJ12hS2_XJ12hS3 | 11 | 1 | 10 |
| XJ12hR1_XJ12hR2_XJ12hR3_vs_XJ24hR1_XJ24hR2_XJ24hR3 | 5 | 2 | 3 |
| XJ12hS1_XJ12hS2_XJ12hS3_vs_XJ24hS1_XJ24hS2_XJ24hS3 | 0 | 0 | 0 |
| XJ24hR1_XJ24hR2_XJ24hR3_vs_XJ24hS1_XJ24hS2_XJ24hS3 | 11 | 5 | 6 |

Note: DEG Set: the name of the differentially expressed gene set; Number of DEGs: the number of differentially expressed circRNAs; Upregulated: the number of upregulated circRNAs; and Downregulated: the number of downregulated lncRNAs.

A hierarchical clustering analysis was performed on the differentially expressed circRNAs that were screened. XJ0hR showed greater differences in differentially expressed circRNAs than XJ0hS (Figure 3).

**Figure 3.**
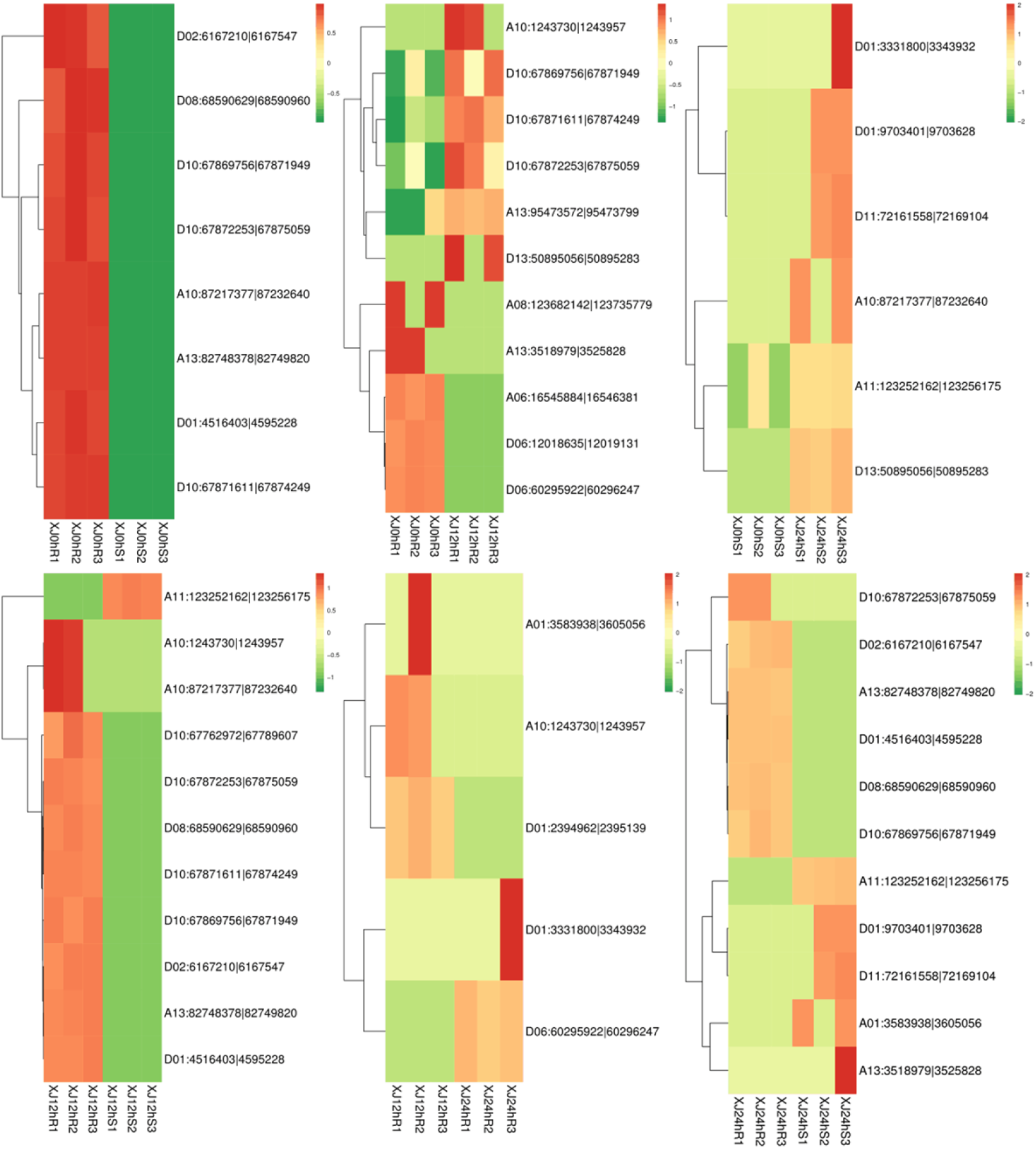
Differentially expressed circRNA expression pattern cluster map Note: Different columns in the figure represent different samples, and different rows represent different circRNAs. The colour represents the expression level of a circRNA in the sample (log10SRPBM+0.000001).

#### 2.3.4 Differential miRNA expression screening

In the process of detecting differentially expressed miRNAs, |log2(FC)|≥0.58 and a P value≤0.05 were used as the screening criteria. Fold change (FC) represents the ratio of expression levels between two samples (groups). The significance p-value obtained from the original hypothesis can be expressed as the probability of expressing no difference. Since the differential expression analysis of miRNAs is an independent statistical hypothesis test for a large number of miRNA expression levels, false positives will be an issue. Therefore, in the analysis process, the Benjamini-Hochberg correction method is sometimes used to test if the original hypothesis is significant. The p-value was corrected, and the false discovery rate (FDR) was finally used as the key indicator for differentially expressed miRNA screening. A total of 453 miRNAs were detected, including 60 known miRNAs and 393 newly predicted miRNAs. The number of differentially expressed miRNAs was small (Table 5).

**Table 5.** Statistics of the number of differentially expressed miRNAs

| DEG Set | Number<br>of<br>DEGs | Upregulated | Downregulated |
| --- | --- | --- | --- |
| XJ0hR1_XJ0hR2_XJ0hR3_vs_XJ0hS1_XJ0hS2_XJ0hS3 | 22 | 8 | 14 |
| XJ0hR1_XJ0hR2_XJ0hR3_vs_XJ12hR1_XJ12hR2_XJ12hR3 | 9 | 3 | 6 |
| XJ0hR1_XJ0hR2_XJ0hR3_vs_XJ24hR1_XJ24hR2_XJ24hR3 | 5 | 3 | 2 |
| XJ0hS1_XJ0hS2_XJ0hS3_vs_XJ12hS1_XJ12hS2_XJ12hS3 | 6 | 3 | 3 |
| XJ0hS1_XJ0hS2_XJ0hS3_vs_XJ24hS1_XJ24hS2_XJ24hS3 | 19 | 13 | 6 |
| XJ12hR1_XJ12hR2_XJ12hR3_vs_XJ12hS1_XJ12hS2_XJ12hS3 | 14 | 7 | 7 |
| XJ12hR1_XJ12hR2_XJ12hR3_vs_XJ24hR1_XJ24hR2_XJ24hR3 | 10 | 7 | 3 |
| XJ12hS1_XJ12hS2_XJ12hS3_vs_XJ24hS1_XJ24hS2_XJ24hS3 | 17 | 7 | 10 |
| XJ24hR1_XJ24hR2_XJ24hR3_vs_XJ24hS1_XJ24hS2_XJ24hS3 | 30 | 15 | 15 |

Note: DEG Set: the name of the differentially expressed gene set; Number of DEGs: the number of differentially expressed miRNAs; Upregulated: the number of upregulated miRNAs; and Downregulated: the number of downregulated miRNAs.

Hierarchical clustering analysis was performed on the selected differentially expressed miRNAs, and the miRNAs with the same or similar expression behaviour were clustered. XJ12hS2 has significantly more differentially expressed miRNAs, and XJ24hR1 had significantly less differentially expressed miRNAs. The clustering results are as follows (Figure 4).

**Figure 4.**
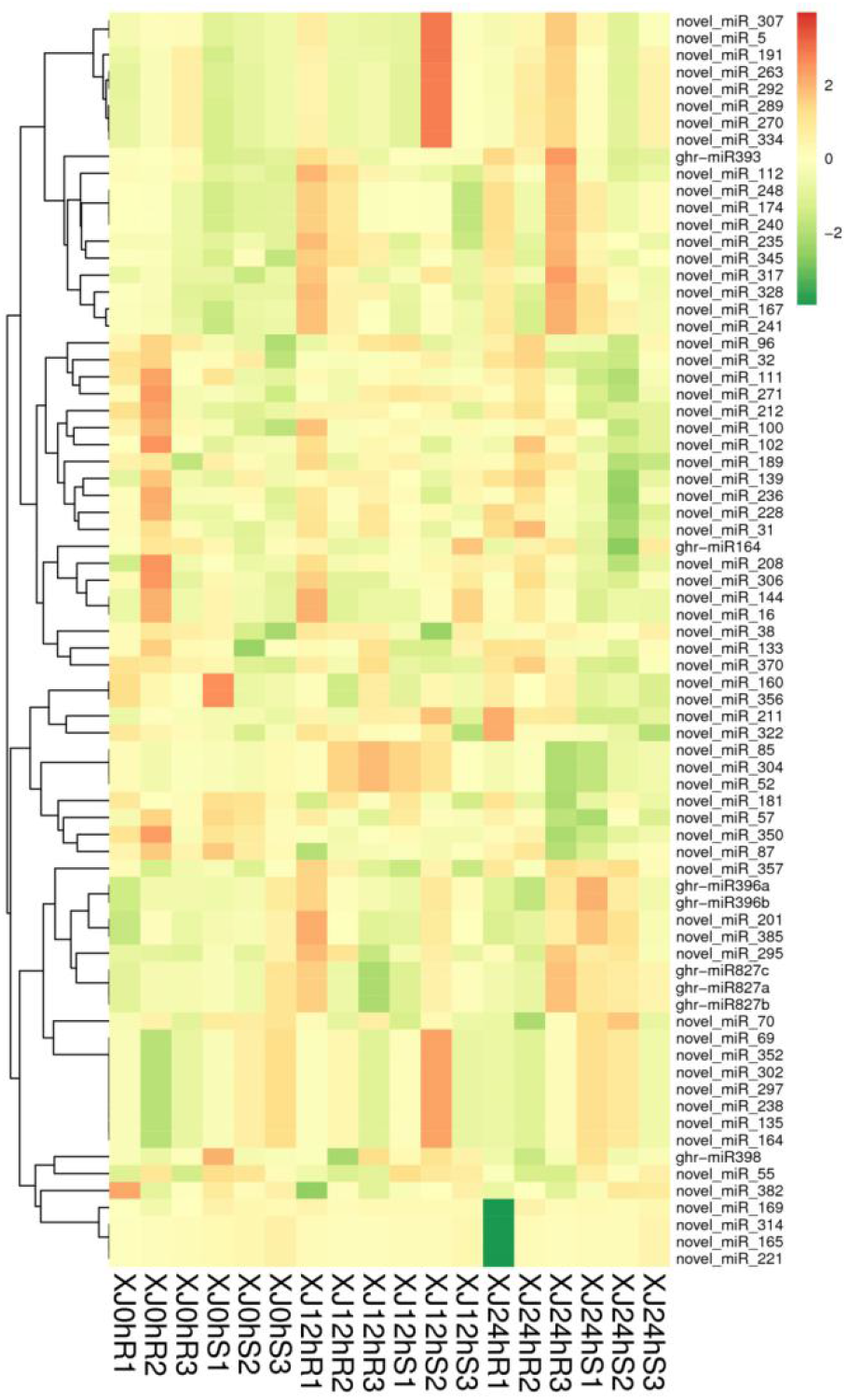
Differentially expressed miRNA cluster map Note: The above figure is a clustering map of differentially expressed miRNAs; the columns represent different samples, and the rows represent different miRNAs. Clustering is based on log10 (TPM+1e^-6^) values. Red represents miRNAs with high expression, and green represents miRNAs with low expression.

#### 2.3.5 qRT-PCR results analysis

Nine genes were randomly selected from the significantly expressed genes for real-time fluorescence quantitative PCR verification. The expression difference between the upregulated genes and downregulated genes was consistent with the trend of the sequencing results, indicating that the sequencing results were reliable (Figure 5).

**Figure 5.**
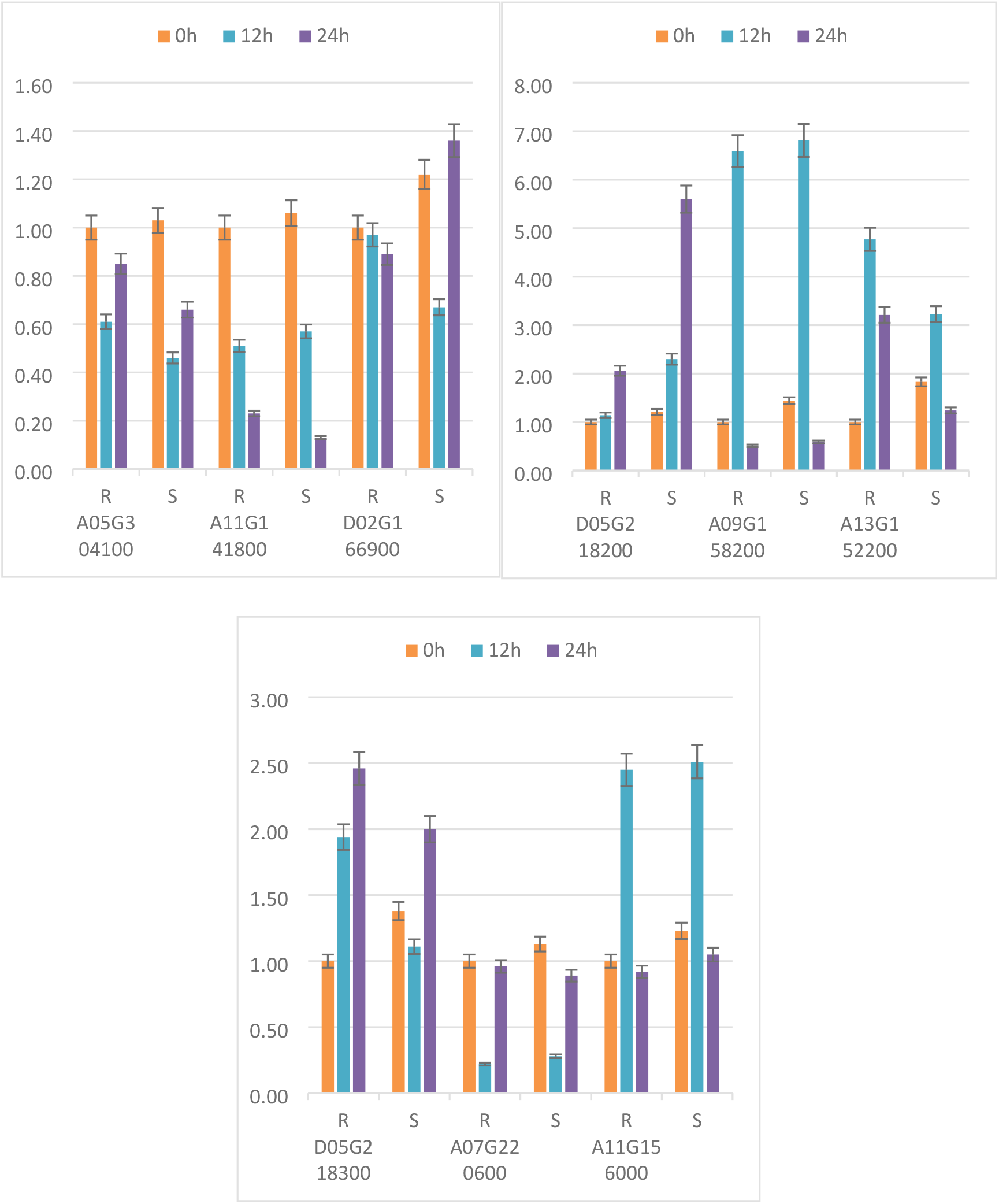
qRT-PCR validation of differentially expressed genes

### 2.4 Functional annotation and enrichment analysis of differentially expressed genes

#### 2.4.1 Functional annotation of differentially expressed genes

The statistics of the number of genes annotated in each differentially expressed gene set are shown in the table below (Table 6).

**Table 6.**
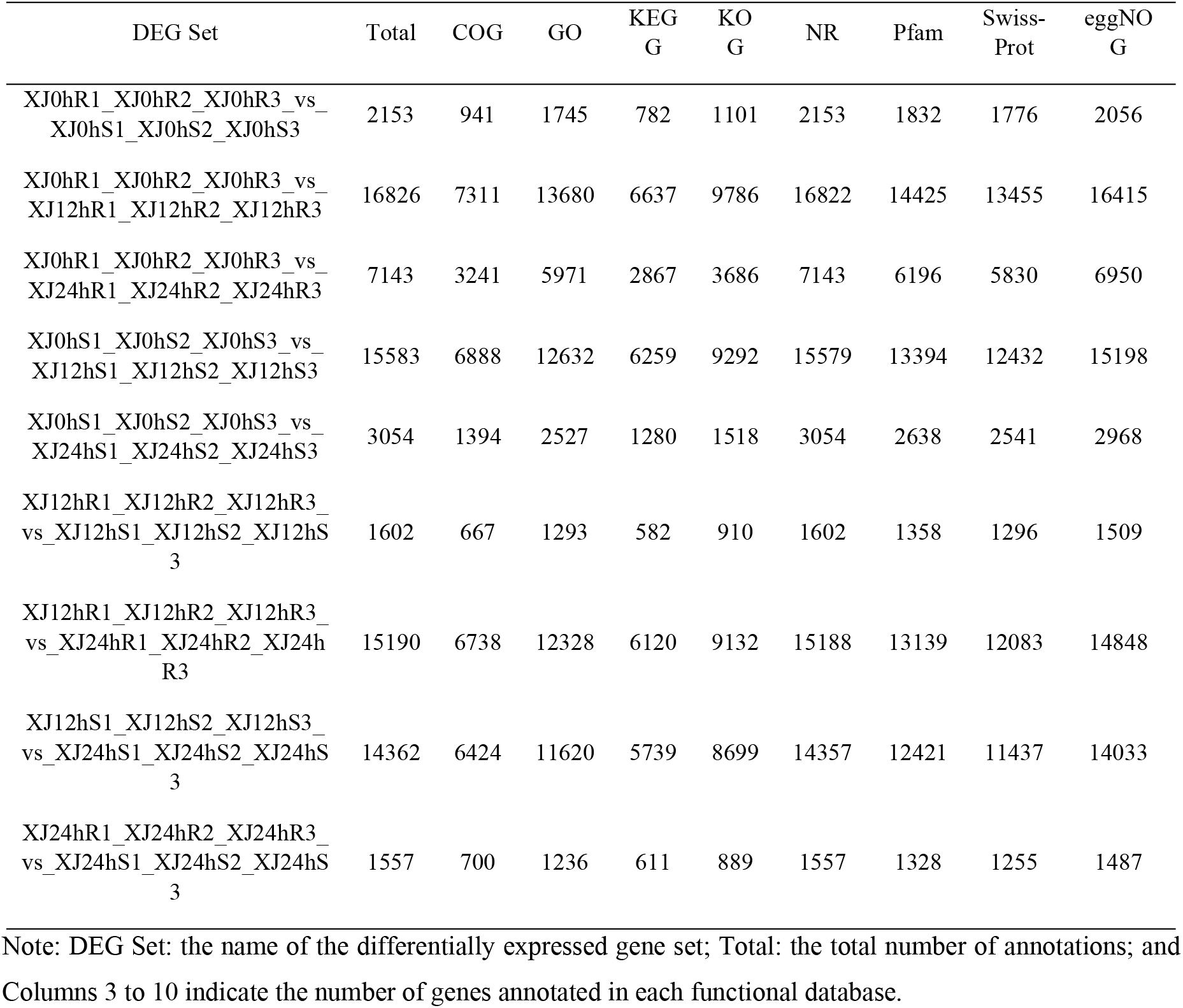
Annotated statistical table of the number of differentially expressed genes

ClusterProfiler (Yu et al. 2012) was used to analyse genes enriched in the biological process, molecular function and cell component categories. Enrichment analysis uses a hypergeometric test method to find GO and KEGG entries that are significantly enriched compared with the entire genomic background. GO entries were significantly enriched in heterocyclic compound binding, and the most enriched functions were classified as response to redox state and pigment binding. The clustering of pathways indirectly demonstrates the importance of certain biological functions. KEGG entries were significantly enriched in signalling pathways such as plant hormone signal transduction. The highly expressed signalling pathway in the 12 h treatment of the two varieties was thiamine metabolism. Thiamine metabolism plays an important role in plant growth and development, biological stress, and nonbiological stress responses. The signalling pathway that was highly expressed in all treatments was glyoxylate and dicarboxylate metabolism (Figure 6), which is consistent with the complexity of the drought stress response.

**Figure 6.**
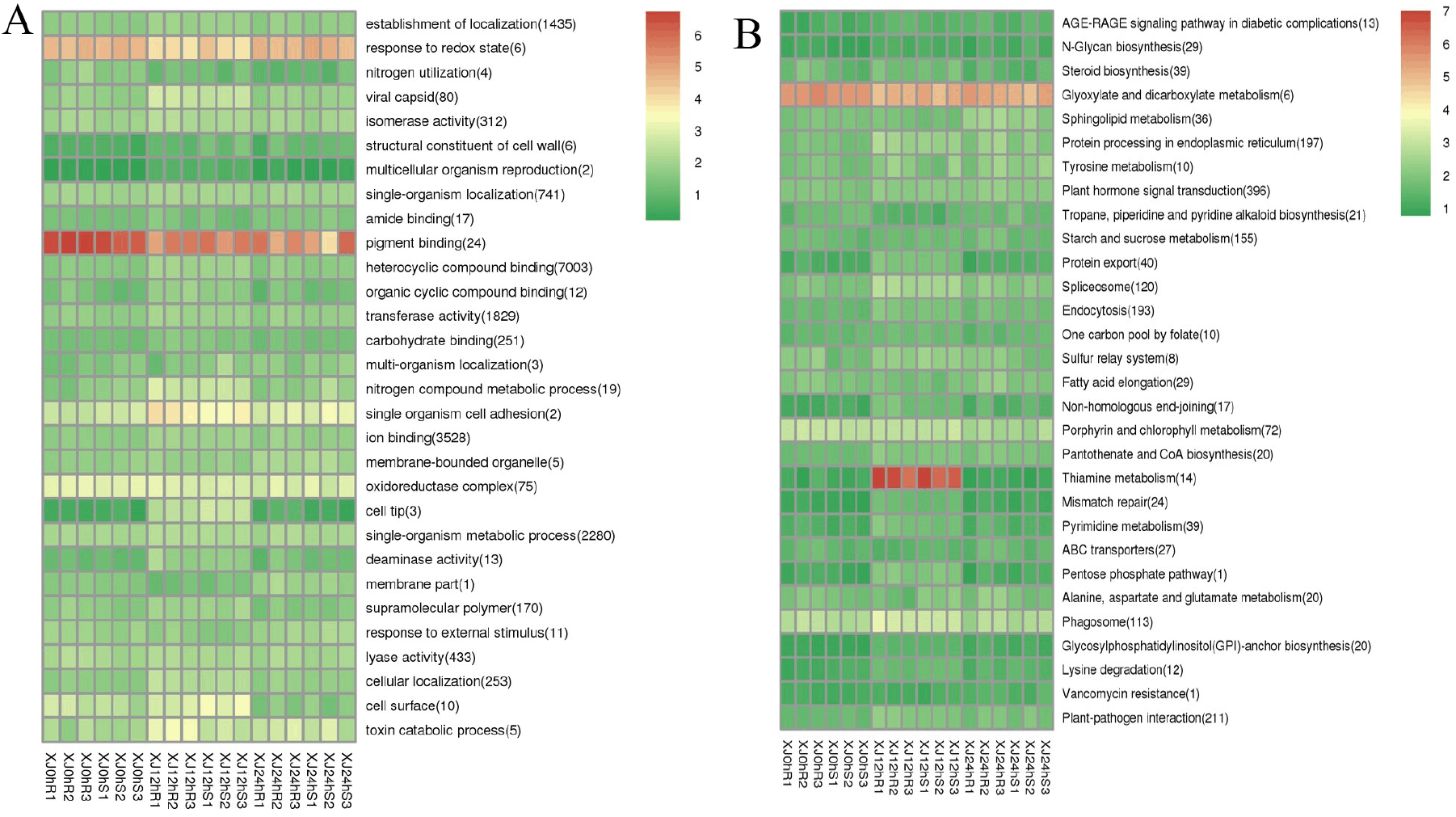
A:GO enrichment cluster map for differentially expressed genes, B:KEGG enrichment cluster map for differentially expressed genes Note: Red indicates metabolic pathways with high expression, and blue indicates metabolic pathways with relatively low expression. The parentheses after the label for each metabolic pathway include the number of genes with significant differences in the metabolic pathway.

#### 2.4.2 Functional annotation of differentially expressed lncRNAs

Differentially expressed lncRNAs were used for GO and KEGG enrichment analysis, and differentially expressed lncRNA target genes were also clustered. The term most significantly enriched with the differentially expressed genes was heterocyclic compound binding, and the most enriched function was classified as organelle membrane and modified amino acid binding (Figure 12).The results showed that the signalling pathway with the most enriched genes was biosynthesis of amino acids, and the highly expressed pathways were photosynthesis, ascorbate and aldarate metabolism, glyoxylate and dicarboxylate metabolism, and inositol phosphate metabolism (Figure 7). The comprehensive analysis of positive and negative effects of LTGs under drought stress revealed the following terms: microtubule-based process (50), organelle membrane (2), nitrogen metabolism (16), cyanoamino acid metabolism (1), and inositol phosphate metabolism (8). These pathways play an important regulatory role in the drought response process.

**Figure 7.**
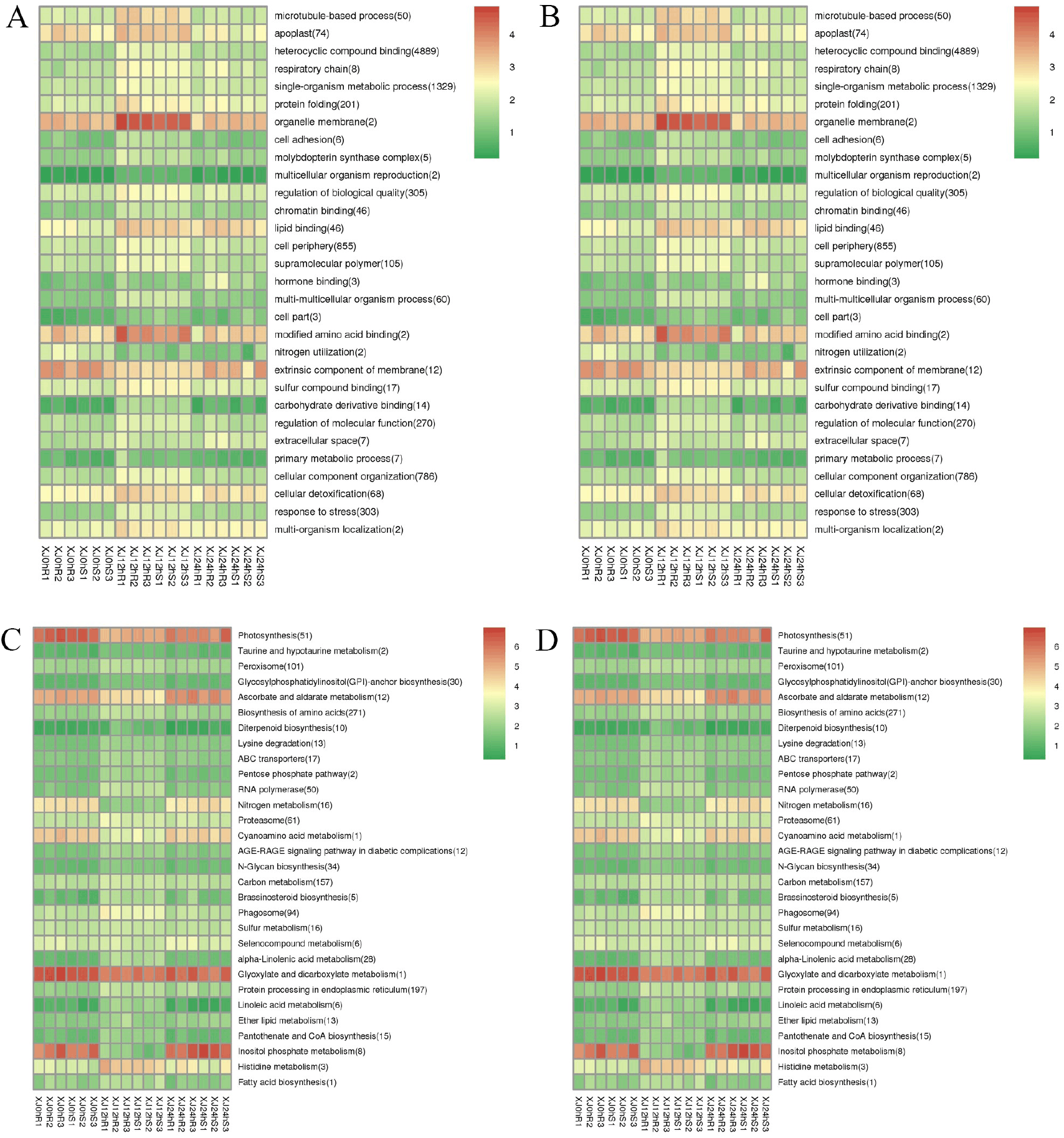
A: Clustering graph of GO enrichment of differentially expressed lncRNA cis-target genes. B: Clustering graph of GO enrichment of differentially expressed lncRNA trans-target genes. C: Cluster map of KEGG enrichment of differentially expressed lncRNA cis-target genes D: Cluster map of KEGG enrichment of differentially expressed lncRNA trans-target genes. Note: Red indicates metabolic pathways with high expression, and blue indicates metabolic pathways with relatively low expression. The parentheses after the label of each metabolic pathway include the number of genes with significant differences in the metabolic pathway.

**Figure 8.**
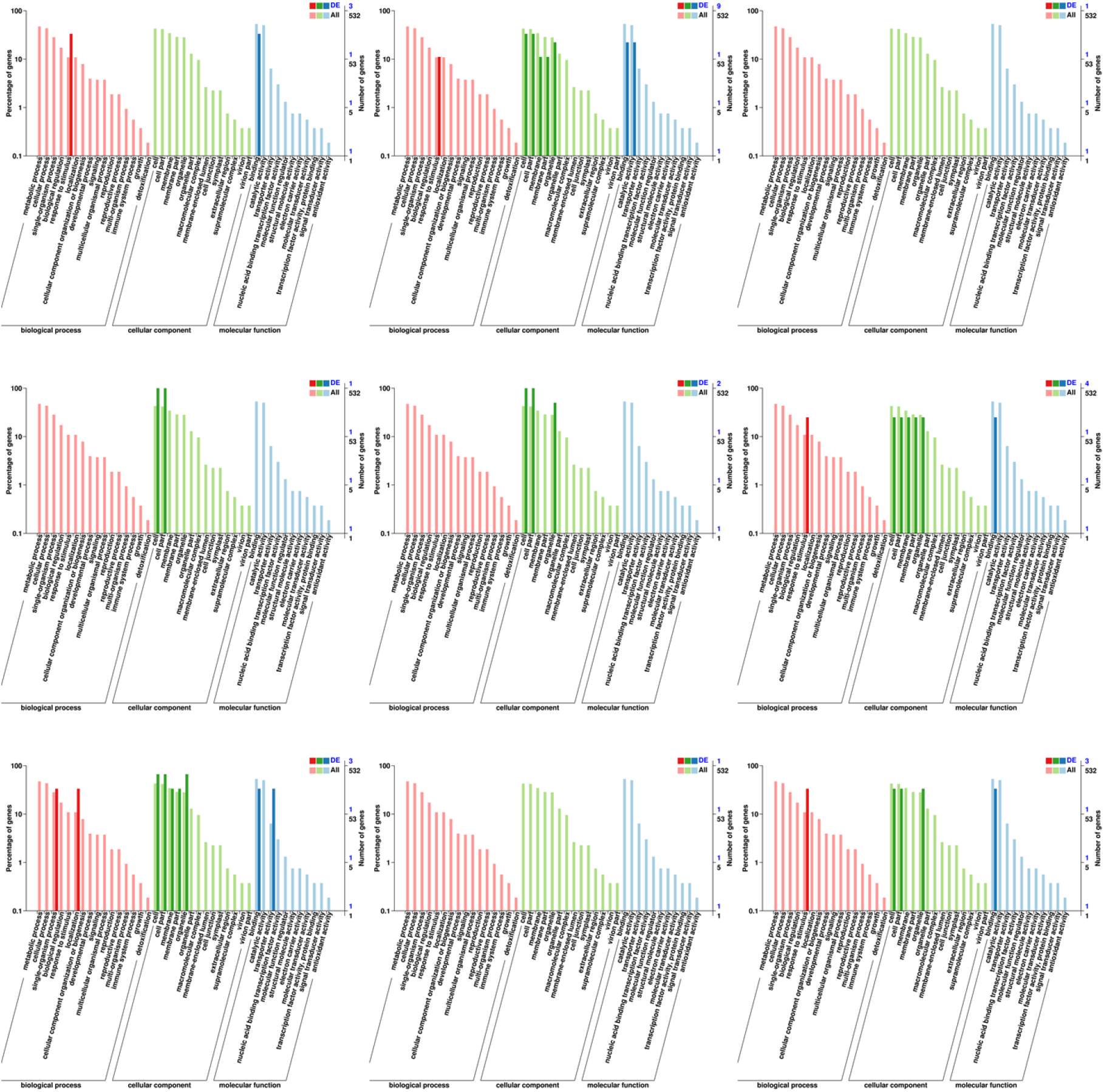
Annotation statistics diagrams of the GO secondary nodes of the different circRNA source genes in each comparison group Note: The abscissa is the GO classification, the left of the ordinate is the percentage of the number of genes, and the right is the number of genes. This figure shows the gene annotation status of each secondary function of GO based on circRNA source genes and all genes. It reflects the status of each secondary function in the two backgrounds. The secondary functions with obvious proportional differences indicate that circRNA source genes are compared with all genes, indicating that if the enrichment trend of genes is different, this function is important.

**Figure 9.**
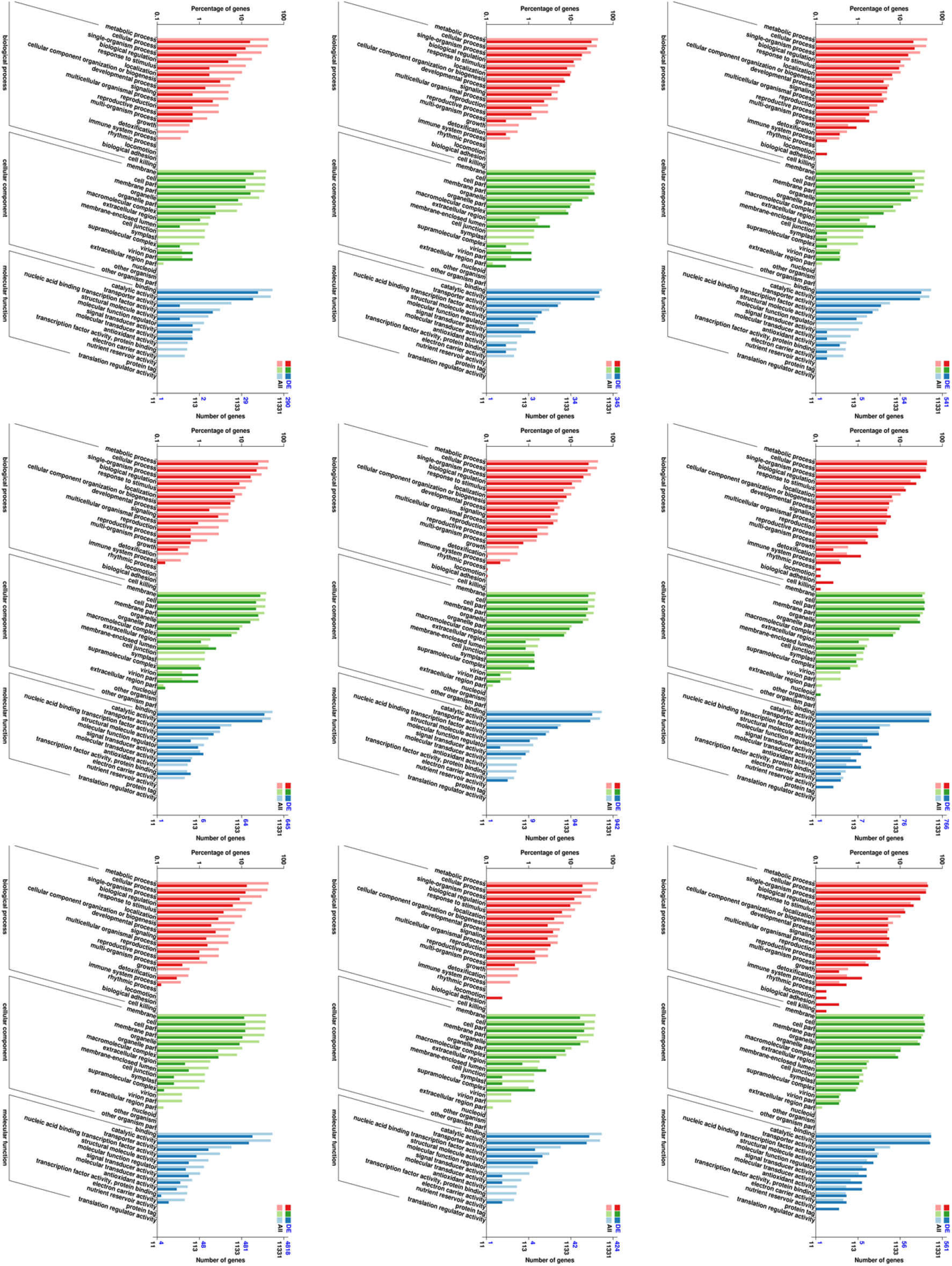
GO annotation classification statistics of differentially expressed miRNA target genes Note: The abscissa is the GO classification, the left of the ordinate is the percentage of the number of genes, and the right is the number of genes. This figure shows the gene enrichment of each secondary function of GO based on the differentially expressed miRNA target genes and the all genes, reflecting the status of each secondary function in the two backgrounds. The secondary functions with obvious proportional differences indicate that circRNA source genes are compared with all genes, indicating that if the enrichment trend of genes is different, this function is important.

**Figure 10.**
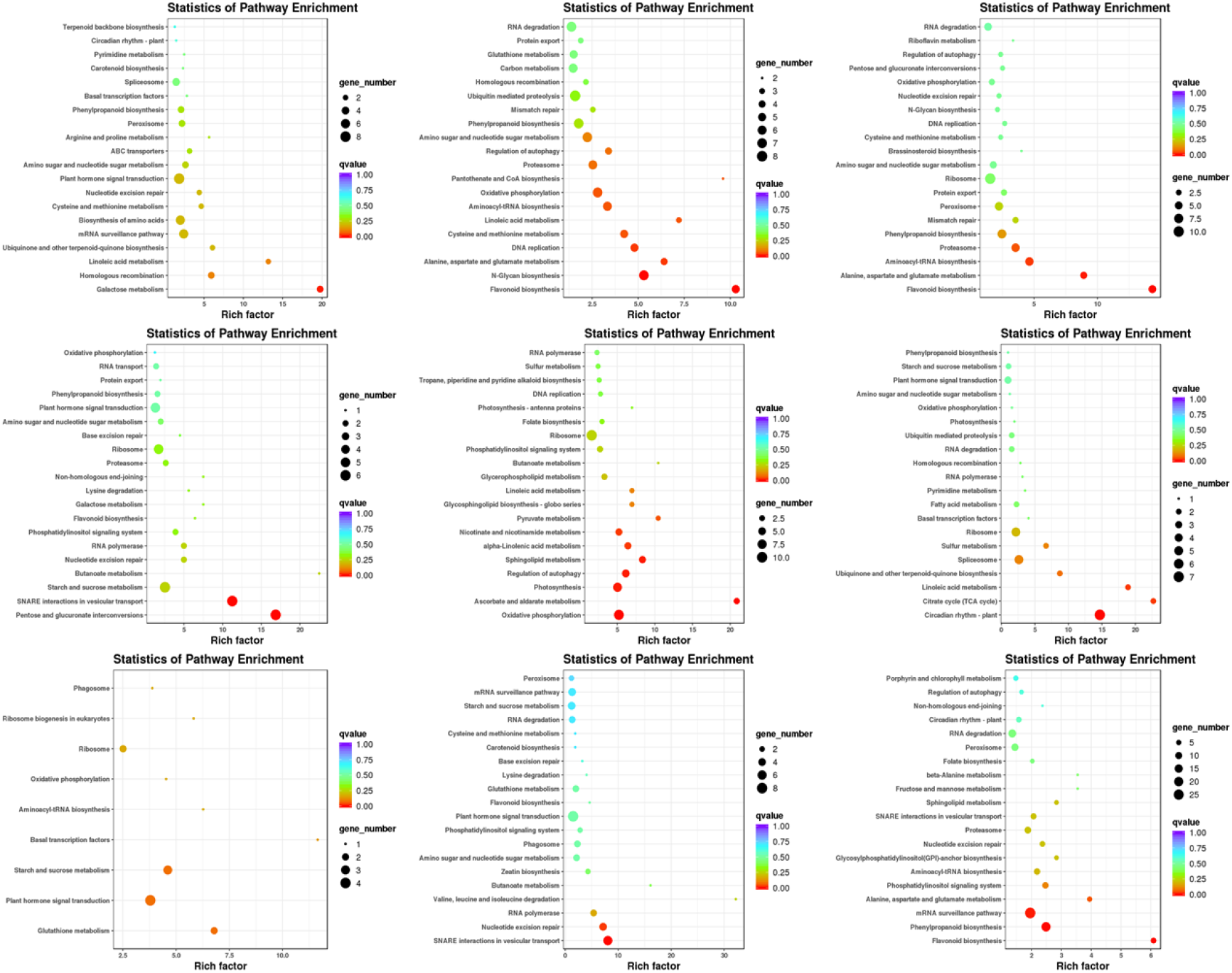
KEGG pathway enrichment scatter diagram of differentially expressed miRNA target genes Note: Each graph in the figure represents a KEGG pathway; the abscissa is the enrichment factor, which represents the proportion of the number of differentially expressed miRNA target genes annotated to a pathway among the total number of genes annotated to the pathway. The larger the value is, the more significant the enrichment level of the differentially expressed miRNA target gene in the pathway. The ordinate is −log10 (Q value), where the Q value is the P value after multiple hypothesis testing correction; therefore, the larger the ordinate is, the higher the enrichment significance of differentially expressed miRNA target genes in this pathway.

**Figure 11.**
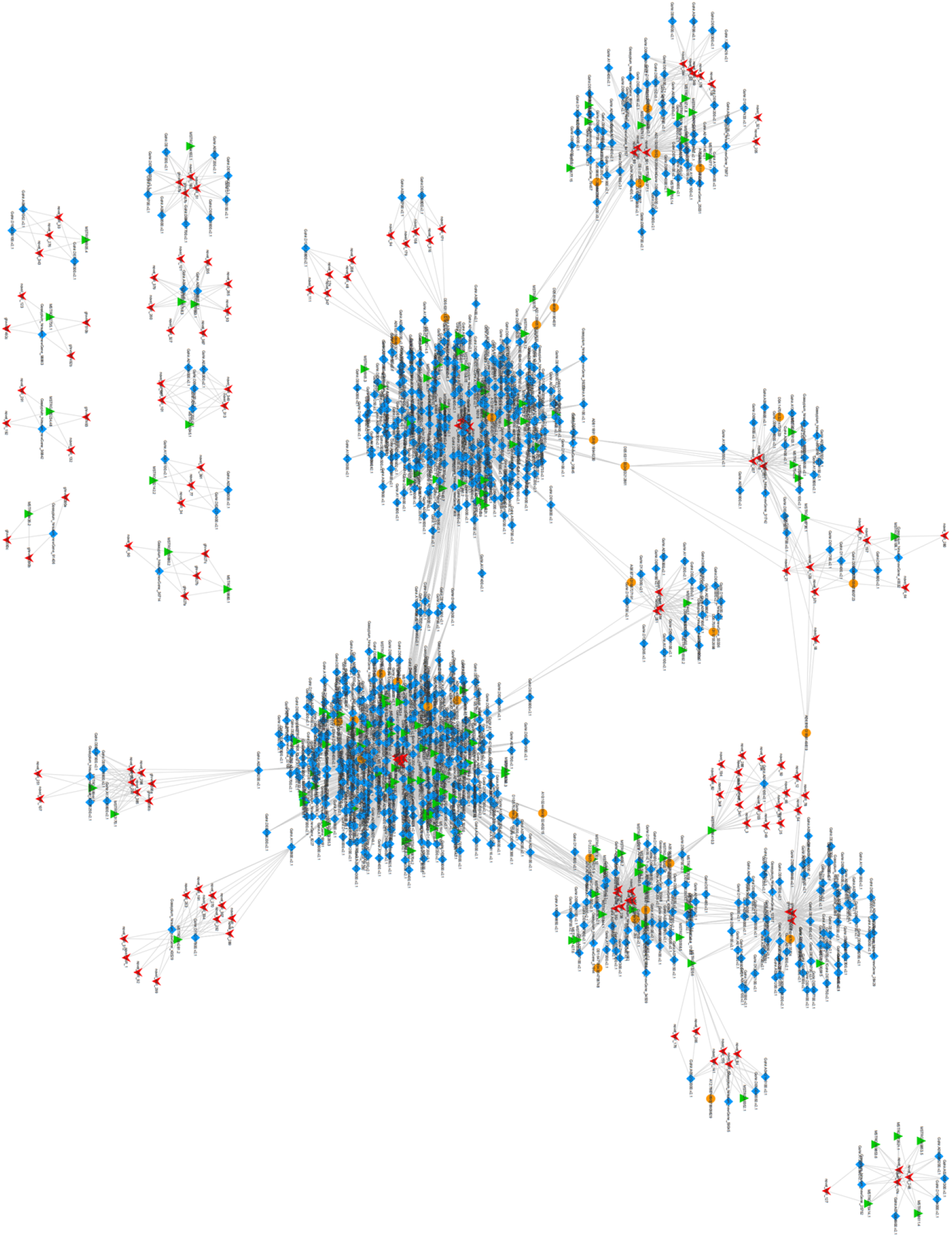
CeRNA regulation network Note: circRNA: round orange. Gene: rhombus blue. lncRNA: Upper triangular green. miRNA: lower arrow red.

**Figure 12.**
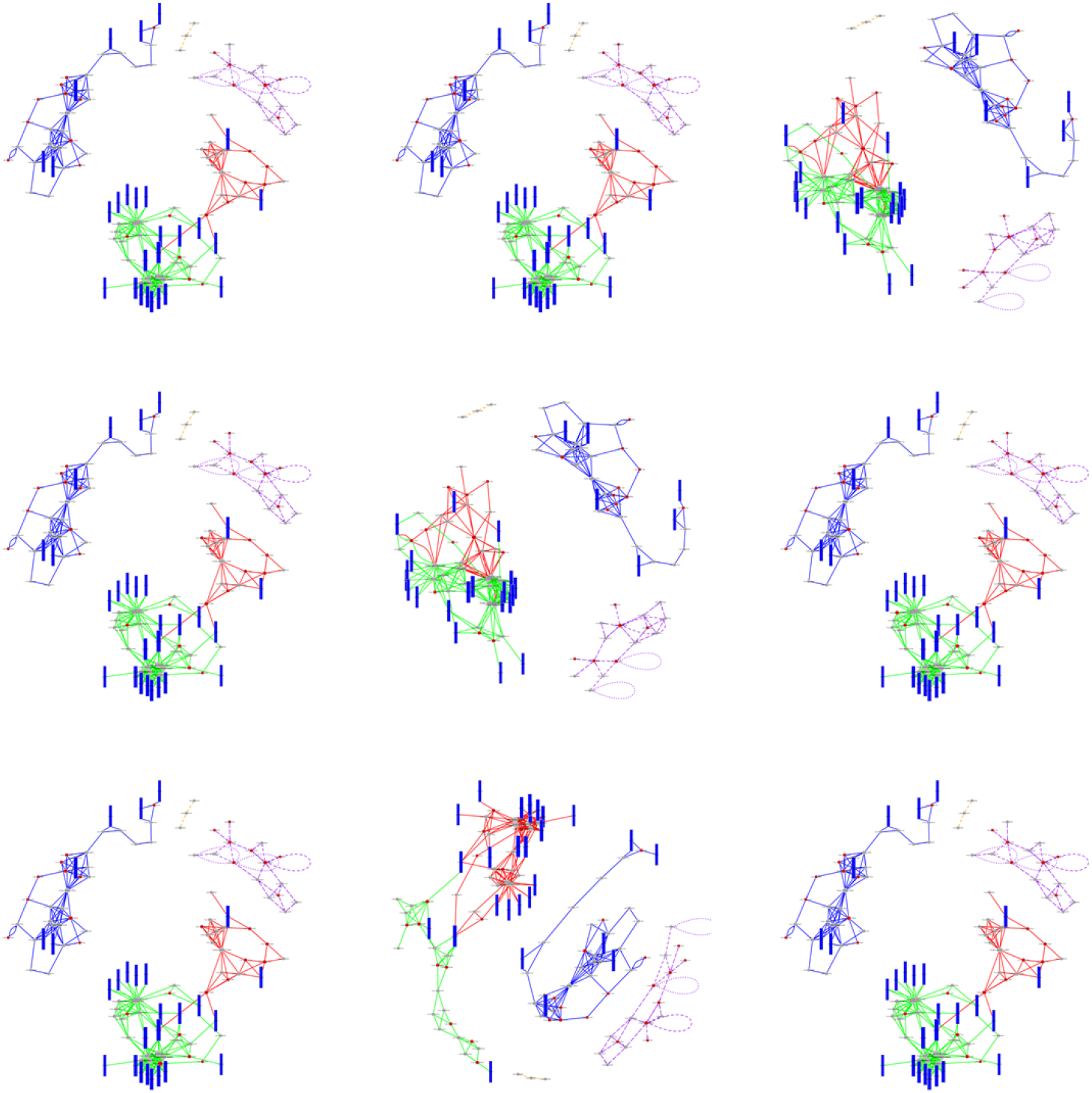
The top 5 most enriched pathways in each comparison group Note: Each dot represents a gene, each rectangle represents a pathway, and the line represents the relationship between genes and between genes and other pathways. The colours of different lines indicate that the relationship comes from different pathways. The red dots are the key genes.

#### 2.4.3 Functional annotation of differentially expressed circRNAs

We performed functional annotations on differentially expressed circRNAs, and the statistical table of the annotation results is shown below (Table 7).

**Table 7.**
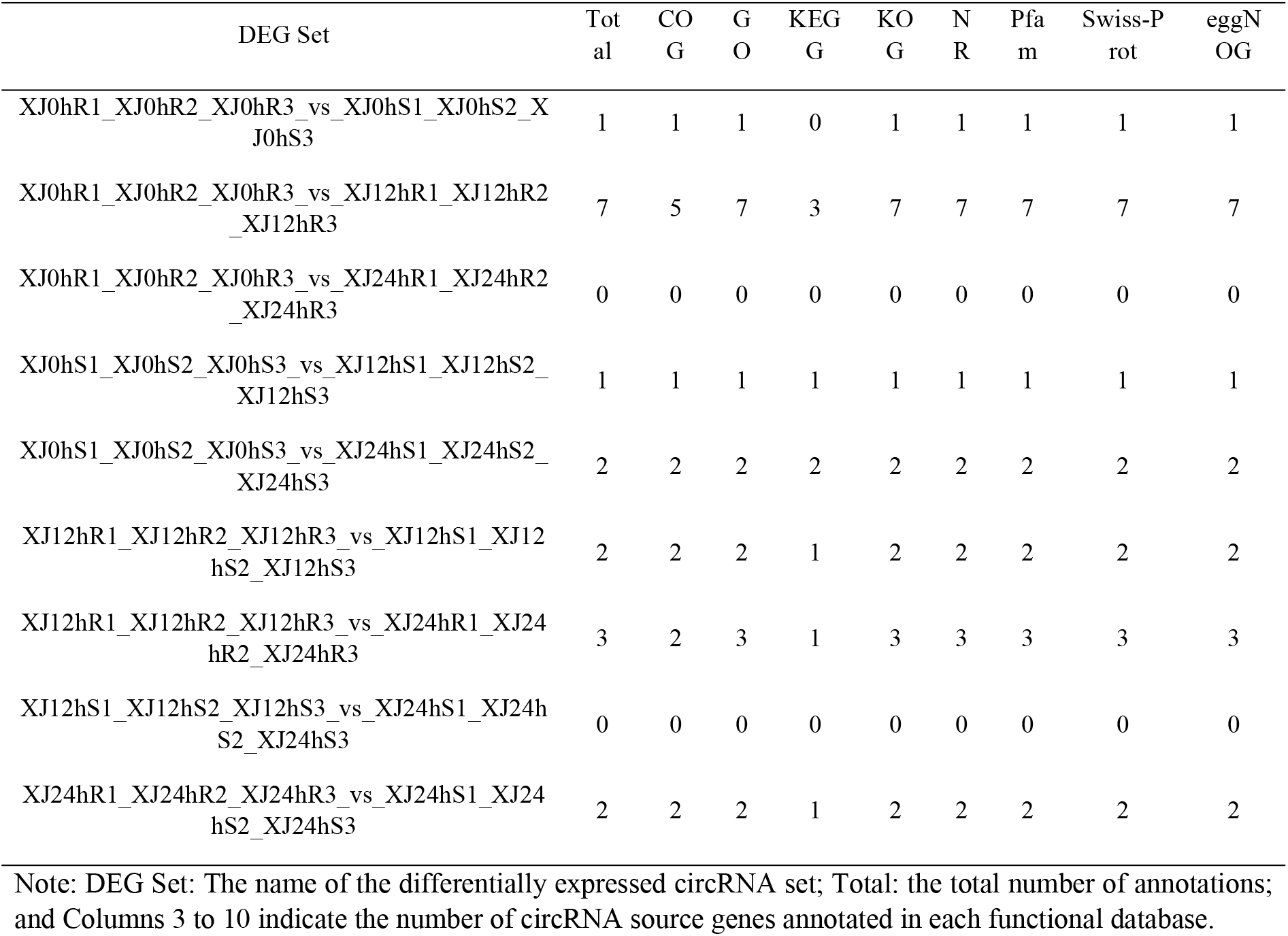
Annotated statistical table of the number of differentially expressed circRNA source genes

GO contains three main categories: biological process, molecular function and cellular component. According to the analysis, the most relevant BP (biological process) terms were response to stimulus and single-organism process, and the most relevant CC (cellular component) terms were cell, cell part, membrane, membrane part and organelle. The most enriched MF ( molecular function) term was related to binding and catalytic activity (Figure 8).

#### 2.4.4 Functional annotation of differentially expressed miRNAs

The statistical results of the number of differentially expressed miRNA target genes annotated between samples are shown in the table below (Table 8).

**Table 8.** Annotated statistics on the number of differential miRNA target genes

| DEG Set | Total | CO<br>G | GO | KE<br>GG | KOG | NR | Pfa<br>m | Swiss-Prot | eggN<br>OG |
| --- | --- | --- | --- | --- | --- | --- | --- | --- | --- |
| XJ0hR1_XJ0hR2_XJ0hR3_vs_<br>XJ0hS1_XJ0hS2_XJ0hS3 | 400 | 156 | 299 | 139 | 223 | 400 | 322 | 304 | 377 |
| XJ0hR1_XJ0hR2_XJ0hR3_vs_<br>XJ12hR1_XJ12hR2_XJ12hR3 | 1009 | 385 | 763 | 377 | 575 | 1009 | 849 | 793 | 949 |
| XJ0hR1_XJ0hR2_XJ0hR3_vs_<br>XJ24hR1_XJ24hR2_XJ24hR3 | 737 | 285 | 559 | 256 | 419 | 737 | 618 | 574 | 690 |
| XJ0hS1_XJ0hS2_XJ0hS3_vs_X<br>J12hS1_XJ12hS2_XJ12hS3 | 376 | 127 | 270 | 119 | 229 | 376 | 310 | 309 | 365 |
| XJ0hS1_XJ0hS2_XJ0hS3_vs_X<br>J24hS1_XJ24hS2_XJ24hS3 | 754 | 269 | 566 | 261 | 439 | 754 | 633 | 589 | 710 |
| XJ12hR1_XJ12hR2_XJ12hR3_v<br>s_XJ12hS1_XJ12hS2_XJ12hS3 | 297 | 109 | 210 | 104 | 181 | 297 | 247 | 226 | 288 |
| XJ12hR1_XJ12hR2_XJ12hR3_v<br>s_XJ24hR1_XJ24hR2_XJ24hR3 | 137 | 46 | 101 | 49 | 75 | 137 | 123 | 118 | 132 |
| XJ12hS1_XJ12hS2_XJ12hS3_vs<br>_XJ24hS1_XJ24hS2_XJ24hS3 | 493 | 185 | 387 | 175 | 321 | 493 | 398 | 404 | 470 |
| XJ24hR1_XJ24hR2_XJ24hR3_v<br>s_XJ24hS1_XJ24hS2_XJ24hS3 | 1999 | 716 | 1530 | 702 | 1175 | 1999 | 1645 | 1547 | 1900 |

Note: DEG Set: The name of the differentially expressed miRNA set; Total: the total number of annotations; and Columns 3 to 10 indicate the number of miRNA source genes annotated in each functional database.

GO enrichment analysis revealed that metabolic process, cellular process and single-organism process were highly enriched in the BP category; membrane, cell, cell part, membrane part and organelle in the CC category; and binding and catalytic activity in the MF category (Figure 9).

KEGG pathway enrichment analysis of the differentially expressed miRNA target genes was performed to determine whether the differentially expressed miRNA target genes were overrepresented in a certain pathway. The enrichment factor was used to analyse the enrichment degree of the pathway, and Fisher’s exact test method was used to calculate the significance of enrichment. The main enrichment pathways were as follows: galactose metabolism; flavonoid biosynthesis; N-glycan biosynthesis; alanine, aspartate and glutamate metabolism; pentose and glucuronate interconversions; SNARE interactions in vesicular transport; oxidative phosphorylation; TCA cycle; circadian plant; and mRNA surveillance pathway(Figure 10).

### 2.5 Construction of the ceRNA regulatory network and integration analysis of key pathways

#### 2.5.1 Construction of the ceRNA regulatory network

CeRNA is a newly identified transcriptional regulatory molecule. CeRNAs can competitively bind to the same miRNA by microRNA response elements (MREs) to regulate their expression levels (Salmena, 2011). The ceRNA hypothesis reveals a new mechanism of RNA interaction. In this study, we used miRNA targeting to obtain candidate ceRNA relationships. A ceRNA relationship network was constructed using Cytoscape, and the ceRNA network contained 1289 edges and 1039 points, including 146 lncRNAs, 859 mRNAs and 34 circRNAs (Figure 11).

#### 2.5.2 Integration analysis of key genetic pathways

Extracting each group of differentially expressed RNAs from the ceRNA relationship pair brings us one step closer in the network, and then the differential ceRNA relationship pair can be obtained (Figure 12). The classic algorithm PageRank in random walk is used to obtain the scores (that is, importance) of all nodes in the network (that is, the different ceRNAs). The key RNA was selected as the key research object if it was ranked in the top 0.05 points in the network. Pathway enrichment analysis was performed on genes in key nodes + lncRNA target genes + circRNA host genes (hereinafter referred to as “key genes”), and the top 5 pathways with the most significant enrichment were selected. The relationship between all genes and genes in these 5 pathways was extracted and integrated into a pathway network, and key genes were mapped to pathways. Here, the node can be regarded as the ID of the ceRNA (gene/lncRNA/circRNA).

#### 2.5.3 Pathway enrichment analysis of critical pathways

Pathway enrichment analysis of the genes in the pathways shows that the top pathways are ribosome biogenesis in eukaryotes; circadian rhythm-plant; N-glycan biosynthesis; pentose phosphate pathway; alanine, aspartate and glutamate metabolism; arginine biosynthesis; folate biosynthesis; 2-oxocarboxylic acid metabolism; glyoxylate and dicarboxylate metabolism; and other glycan degradation (Figure 13).

**Figure 13.**
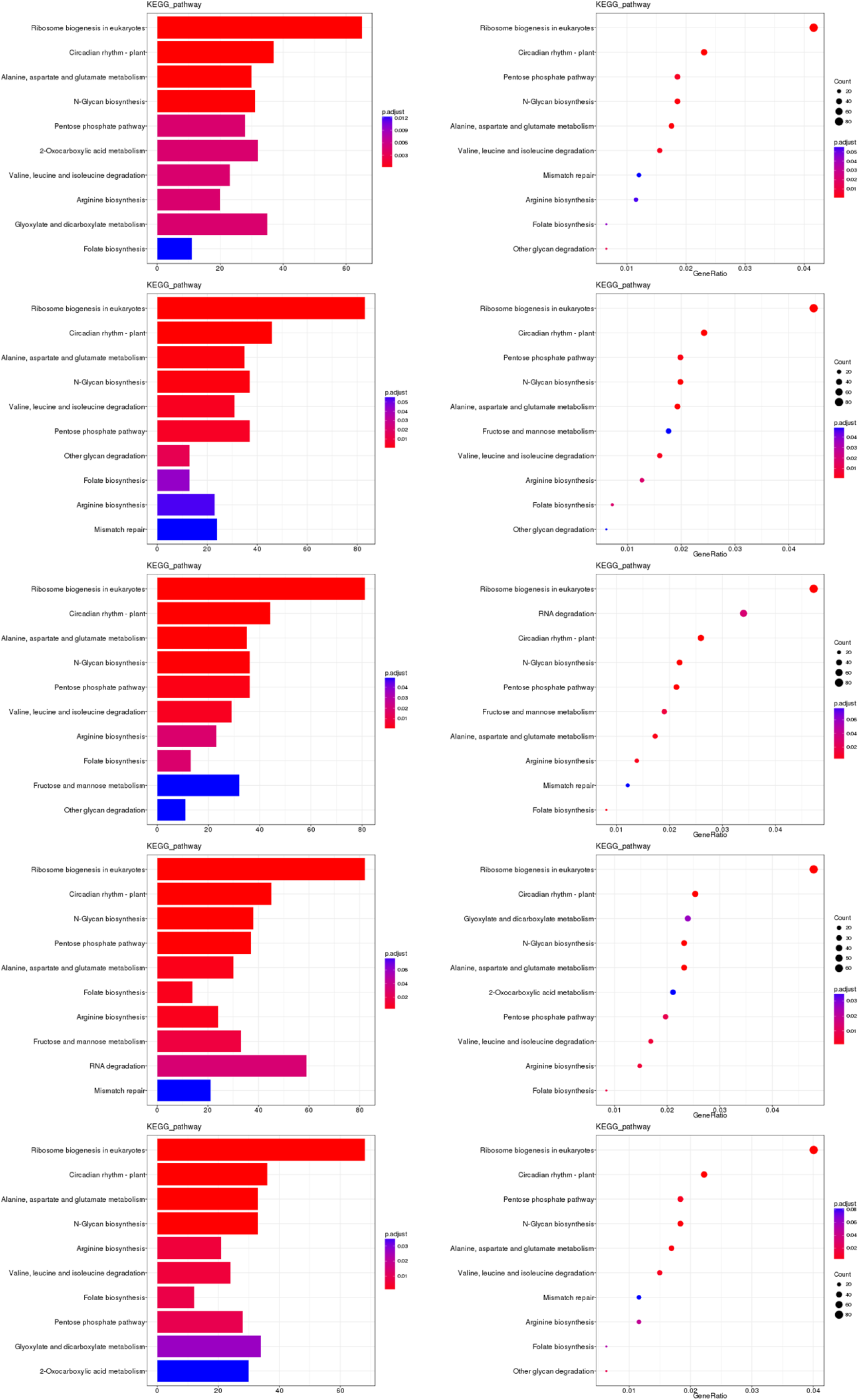
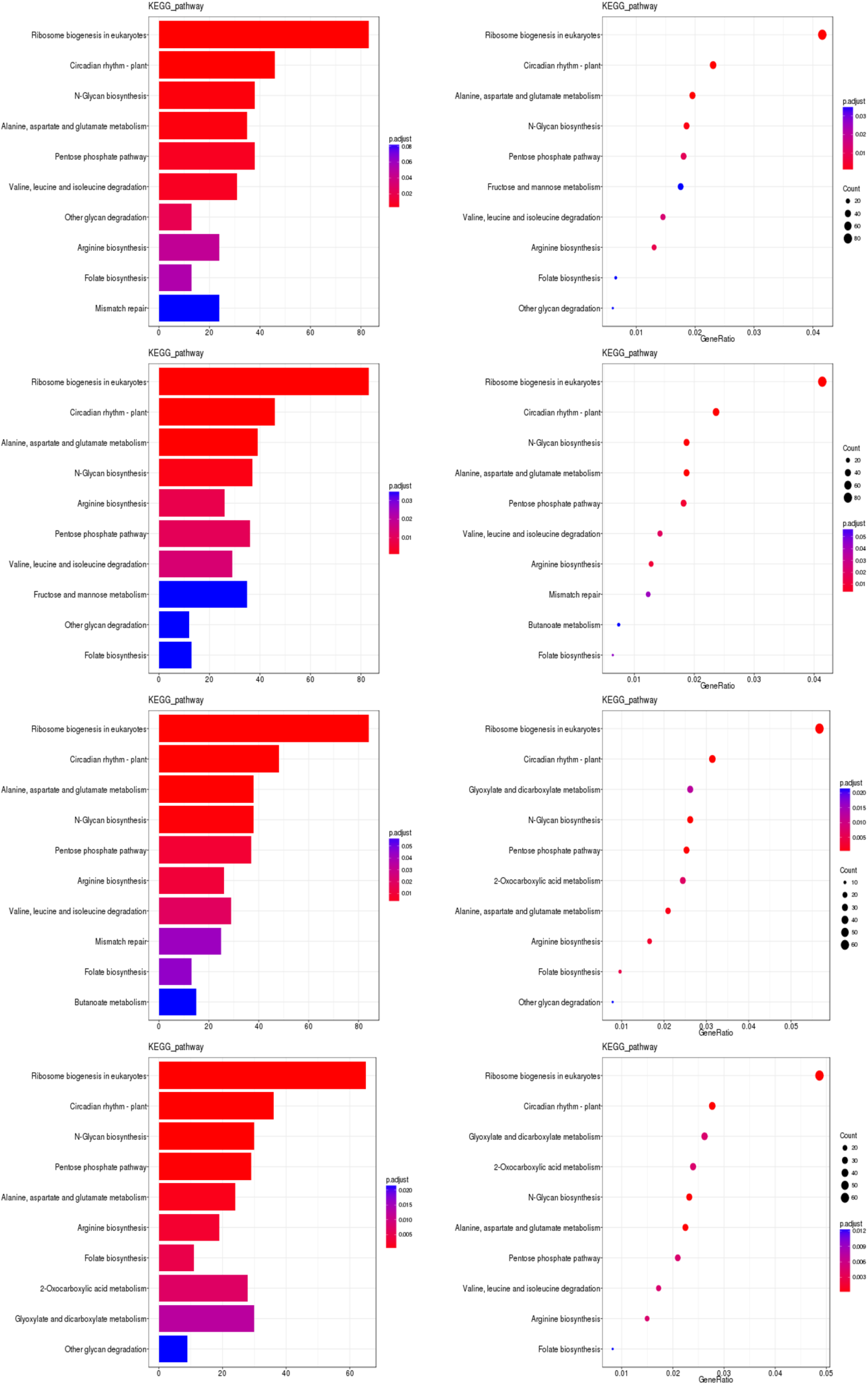
Bar chart and bubble chart of KEGG enrichment of key genes in each comparison group Note: KEGG enrichment bar graph: the abscissa is GeneNum, which is the number of genes of interest annotated in the entry, and the ordinate is each pathway entry. The colour of the column represents the p-value of the hypergeometric test. KEGG enrichment bubble chart: the abscissa is GeneRatio, which is the ratio of the gene of interest annotated in this entry to the number of differentially expressed genes, and the ordinate is each pathway entry. The size of the dot represents the number of differentially expressed genes annotated in the pathway, and the colour of the dot represents the p-value of the hypergeometric test.

### 2.6 Differential gene annotation and ceRNA network analysis were used to screen drought resistance-related genes

Through the integration analysis of key genes in the key pathways and the annotation analysis of GO and KEGG, combined with previous research progress, it was found that there may be drought-related genes that may be key genes in the alanine, aspartate and glutamate metabolism pathway (ko00250) (Table 9). Two genes, Gohir.A11G156000.v2.1 and Gohir.A07G220600.v2.1, were annotated and screened. Compared with the 0 h treatment, in the 12 h treatment, the A11G15600 gene was upregulated and the A07G220600 gene was downregulated in the same variety. Compared with the 12 h treatment, in the 24 h treatment, the A11G15600 gene was downregulated and the A07G220600 gene was upregulated. Through GO and KEGG analysis, the functions of these two genes were identified. Coexpression trend analysis of the selected drought resistance genes provided a more intuitive understanding of the expression trends of these genes between samples. Gohir.A11G156000 was upregulated at 0 vs. 12 h and downregulated at 12 vs. 24 h. Gohir.A07G220600 was downregulated at 0 vs. 12 h and upregulated at 12 vs. 24 h. The comparison of Gohir.A11G156000.v2.1 with Tair3 shows that the homologous gene in *Arabidopsis thaliana* is AT3G22200, with 77% homology, and the gene is POP2, a pyridoxal phosphate (PLP)-dependent transferase superfamily protein. The homologous gene for Gohir.A07G220600.v2.1 in *Arabidopsis thaliana* is AT5G14760, with 76% homology, which encodes an AO-L-aspartate oxidase (Figure 14).

**Figure 14.**
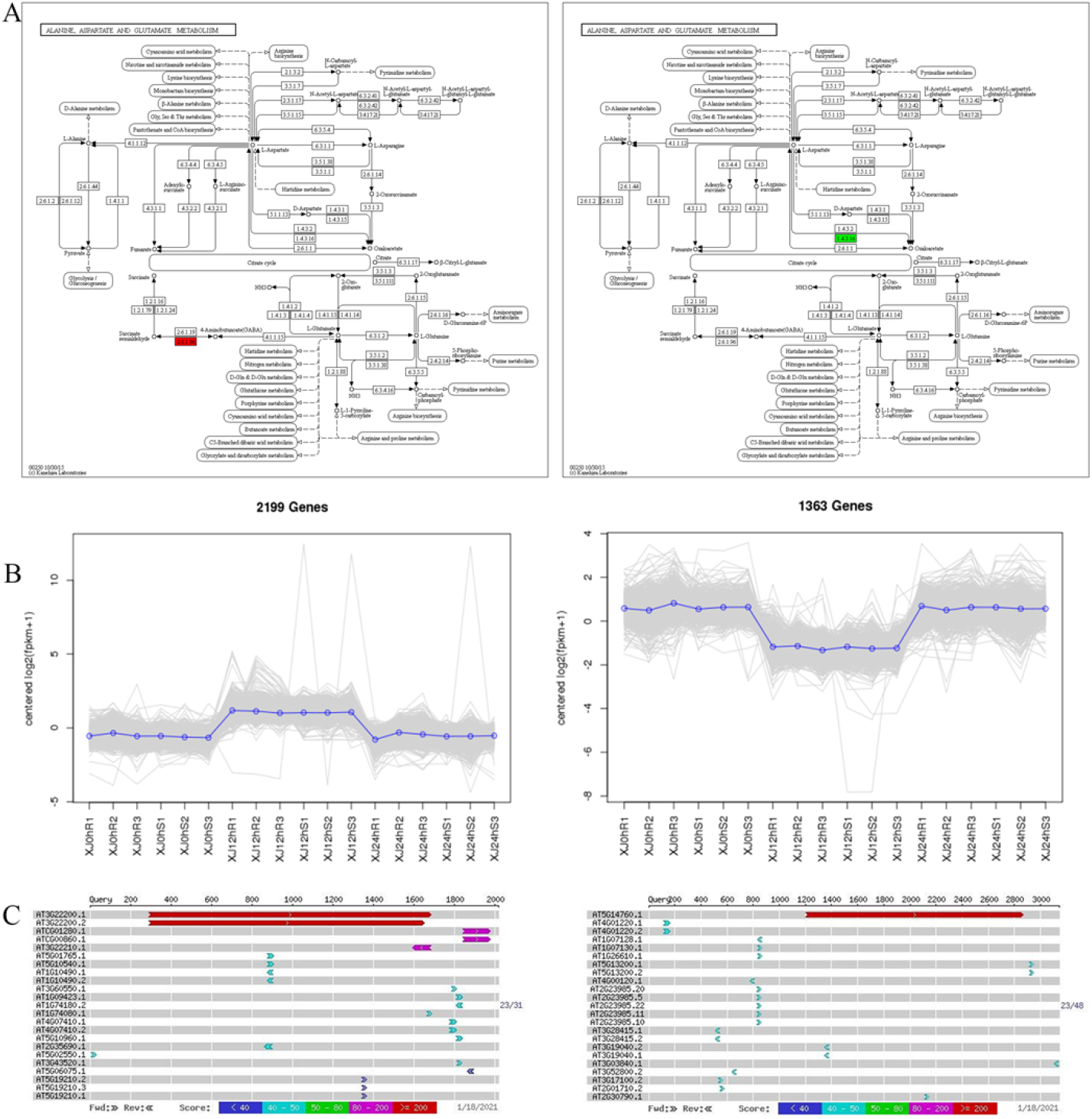
Drought-resistant gene screening. A: KEGG annotation path diagram. The number in the box represents the number of enzymes (EC number), and the entire pathway is composed of a complex biochemical reaction catalysed by a variety of enzymes. In this pathway diagram, the enzymes related to differentially expressed genes are marked with different colours. B: Gene coexpression trend graph. The abscissa represents the sample or time point, and the ordinate represents logarithmic centred expression levels. The grey line in the figure represents the expression trend of a gene. The blue line indicates the type of gene. C: Gene sequence blast results.

**Table 9.** Alanine, aspartate and glutamate metabolism pathway key regulatory genes

| Key gene | Homologous genes in *Arabidopsis* |
| --- | --- |
| Gohir.A11G156000 | AT3G22200 |
| Gohir.A11G227533 | AT3G27740 |
| Gohir.D01G114300 | AT5G10920 |
| Gohir.D04G022000 | AT3G24090 |
| Gohir.D05G088300 | AT3G47340 |
| Gohir.A07G220600 | AT5G14760 |

## 3 Discussion

### 3.1 Rapid screening of drought resistance-related genes by differential gene annotation and ceRNA regulatory network analysis

A ceRNA can be used as a bait to attract an miRNA and isolate it, which can relieve the inhibition of target genes by the miRNA. With MRE as the communication language, mRNA, lncRNA, circRNA and pseudogenes can achieve mutual regulation through miRNA competition mechanisms, forming a large-scale posttranscriptional regulatory network called a ceRNA network.

Franco-Zorrilla et al. (2007) first discovered noncoding RNAs with miRNA bait functions in Arabidopsis. Under low phosphorus stress, lncRNA IPS1 competed with PH02 to bind miR399. Whole transcriptome sequencing (Deng et al., 2018) of cotton trifoliate seedlings under salt stress revealed 44 lncRNAs and 6 miRNAs. Differential expression was verified by qRT-PCR, in which lnc973 and lnc253 competed to bind miR399 and miR156e, respectively.

Although ceRNA was first found in plants, its research progress was far behind that in humans and mammals. The number of ceRNA-related studies included in PubMed continued to grow from 2011 to 2018, but the number of plant-related ceRNA reports was relatively small. The proportion of plant-related reports in 688 articles in 2018 was only 9.74%.

In this experiment, the genes obtained were combined with the cotton genome database through functional annotation of differential genes and ceRNA network screening, and finally, the key genes were identified. This is a rapid method to obtain drought-related genes. Through the construction of a ceRNA network, more critical regulatory genes that can directly act on and help plants respond to drought stress signals were identified,. Finally, two drought-related candidate genes with high expression under drought stress were found.

### 3.2 Study on GABA in plant stress resistance

The A11G156000 homologous gene was predicted to be the POP2 gene, encoding GABA-T. GABA aminotransferase (GABA-T) catalyses the conversion of GABA in the GABA shunt pathway to hemisuccinic acid (SSA), and some studies have shown that it is related to plant stress resistance. In plants, GAD (glutamic acid decarboxylase) present in the cytoplasm and GABA-T and SSADH (succinic semialdehyde dehydrogenase) present in mitochondria jointly regulate GABA branch metabolism, where GAD is the rate-limiting enzyme for the synthesis of GABA. γ-Aminobutyric acid (GABA) is a four-carbon, nonprotein amino acid, and its chemical formula is C4H9NO2 (GABA). Since it was first found in potato tubers in the 1950s, it has attracted wide attention. Subsequent studies have found that GABA widely exists in vertebrates, plants and microorganisms. GABA is produced in the cytoplasm (Shelp et al, 1995). The content of GABA in plant tissues is very low, usually between 0.3 and 32.5 μmol/g. It has been reported that the accumulation of GABA in plants is related to the stress response, which can lead to the rapid accumulation of GABA under hypoxia, heat, cold, mechanical damage and salt stress. In higher plants, the metabolism of GABA is mainly completed by three enzymes. First, under the action of GAD, the irreversible decarboxylation of L-glutamic acid at the α-position occurs, and the released products GABA, PLP (pyridoxal 5-phosphate monohydrate) and GAD recover to their initial state at the end of the reaction. Then, under the catalysis of GABA-T (GABA transaminase), GABA reacts with pyruvate and α-ketoglutaric acid to form succinic semialdehyde. Finally, SSADH catalyses the oxidative dehydrogenation of succinic semialdehyde to form succinic acid and finally enters the tricarboxylic acid cycle (Krebs circle) (Wang et al. 2018; BOUCHE et al.,2003). This metabolic pathway forms a branch of the TCA cycle called the GABA shunt. Previous studies have shown that changes in TCA cycle enzyme activity revealed the metabolic function of GABA and have demonstrated the close relationship between GABA and respiration (Palanivelu R et al, 2003; Renault H et al, 2011). GABA also acts as a regulator of oxidative metabolites. The SSADH mutant of *Arabidopsis thaliana* was exposed to high temperature, and the accumulation of reactive oxygen intermediates (ROIs) was found, resulting in plant death, which showedthat there was a relationship between the ROIs and GABA. Similarly, SSADH and GABA-T gene mutants have a large number of ROIs at high temperature. The use of the ROI elimination agent PBN can cause GABA accumulation, thereby improving the survival rate of yeast. Therefore, it is believed that the GABA shunt process can reduce the accumulation of ROIs to protect organisms from oxidative damage and oxidative decay caused by stress.

Based on the above analysis, the following inferences can be made. In the alanine, aspartate and metabolic pathways, the A11G156000 gene was upregulated at 0h vs 12 h, indicating that GABA-T rapidly accumulated in cotton under drought conditions. GABA could reduce the damage to stressed plants by reactive oxygen species and improve the activity of protective enzymes in plants. The 12h vs 24 h downregulation indicated that PLP and GAD returned to their initial state at the end of the reaction. Under drought conditions, the growth and leaf area extension of roots and stems were inhibited, and the levels of reactive oxygen species increased. The production of low molecular osmotic adjustment substances such as GABA and other amino acids, polyols and organic acids increased, and the expression of enzymes involved in antioxidant injury was upregulated (FAROOQ et al.,2009).

### 3.3 Mechanism of L-aspartate oxidase and NAD in plant stress resistance

The homology comparison results within *A. thaliana* predicted that the homologous sequence of AO wasA07G220600 in *A. thaliana*, which is an L-aspartate oxidase encoding the early steps of NAD (nicotinamide adenine dinucleotide) biosynthesis (Rongvaux et al., 2003). NAD is a ubiquitous coenzyme involved in redox reactions and can be converted between oxidated and reduced forms without any net consumption. NAD is mainly converted into its reduced form NADH in a catabolic reaction, and NADH is mainly oxidized through the mitochondrial electron transfer chain and participates in the redox process of cells. NADH can react with free radicals to inhibit peroxidation. Arabidopsis synthesizes NAD from Asp using AO, QS, and QPT, each of which is essential for plant growth and development. These enzymes are found in the plastid, suggesting that the early steps in the biosynthesis of NAD occur in this organelle (Akira Katoh, et al, 2006). The equation of Asp catalysed by L-Asp oxidase is as follows: L-aspartic acid + O2 = iminoaspartic acid (α iminosuccinic acid) + H2O2. The results showed that L-Asp oxidase was related to the increase in reactive oxygen species (Katoh and Hashimoto, 2004).

Based on the above analysis, we can make the following inferences: In the alanine, aspartate and glutamate metabolism pathways, the A07G220600 gene was downregulated in 0h vs 12 h, indicating the stomatal closure of cotton under drought conditions, which weakened respiration and reduced the production of reactive oxygen species to improve stress resistance. The upregulation of 12h vs 24 h indicated that under drought stress, reactive oxygen species in plants increased significantly. L-Aspartate oxidase catalysed L-aspartate binding to oxygen and caused a ROS burst. The increase in L-aspartate oxidase expression increased NADH synthesis to improve plant antioxidant capacity. Under drought conditions, the growth and leaf area extension of roots and stems were inhibited, and reactive oxygen species increased. The production of low molecular weight osmotic adjustment substances such as GABA and other amino acids, polyols and organic acids increased, and the expression of enzymes involved in antioxidant injury was upregulated.

Studies have shown that under drought conditions, the expression of genes related to intracellular homeostasis, active oxygen scavenging, structural protein stability protection, osmotic regulators, transporters and so on is upregulated. In this experiment, highly expressed genes were involved in regulating a variety of drought resistance-related pathways, and they worked together to improve the drought resistance of plants. At the same time, the results also confirmed the effectiveness of the expression regulatory network, which could be used to explore more multifunctional genes.

## 4. Conclusion

This study is the first to use three-leaf cotton plants for hydroponic experiments with 17% PEG solution for simulating drought stress. We analysed the transcription levels of all genes in 0 h, 12 h, and 24 h leaf samples from different varieties and different treatment timepoints. GO clustering of differentially expressed genes, KEGG clustering, pathway analysis and the construction of ceRNA networks, along with the synthesis of existing literature reports, allowed us to screen for drought resistance-related pathways in cotton, key genes in KEGG pathways, and candidate salt resistance genes. The main conclusions are as follows:

The KEGG analysis of the differentially expressed genes show enrichment in metabolic pathways such as plant hormone signal transduction, carbon metabolism, and photosynthesis-related pathways, indicating that these processes may play an important role in plant growth and development and in resisting drought stress. Hormones in the phytohormonal signalling pathway regulate each other and can be used as secondary signalling molecules to regulate the expression of downstream stress-related genes. Carbohydrates can not only be used as an energy source but can also be used as signalling molecules to regulate plant growth and resist adversity. Photosynthesis-related pathways are also significantly enriched. Photosynthesis-produced organic matter can provide energy for plant growth and development. Cotton can induce a large number of functional proteins to adjust to stressful environments and to improve resistance to stress. Under drought stress, cotton reduces oxidative damage by synthesizing a large number of enzymes associated with ROS removal.

Two drought-related candidate genes (Gohir.A11G156000, Gohir.A07G220600) were obtained. Subsequent genetic function verification can be carried out, and the reliability of drought-resistant genes can be quickly screened using ceRNA regulatory networks.

## 5. Materials and Methods

### 5.1. Plant materials and experimental reagents

The upland cotton (*Gossypium hirsutum* L.) varieties used in this experiment are the drought-resistant material Xinluzhong No. 82 and the drought-sensitive material Kexin No. 1. The materials were provided by the Upland Cotton Research Group of the Economic Crop Research Institute of Xinjiang Academy of Agricultural Sciences.

Hoagland solution preparation was purchased from Baierdi Biotechnology Co., Ltd. (Beijing, China), and the RNAprep Pure polysaccharide polyphenol plant total RNA extraction kit, Fasting RT Kit (with gDNase), and SuperReal PreMix Plus (SYBR Green) were purchased from TIANGEN Co., Ltd. (Beijing, China).

The primers were synthesized by Beijing Bomai Biotechnology Company.

### 5.2 Experimental method

#### 5.2.1 Setting of processing conditions and extraction of total RNA

Filter paper was used to germinate seeds for 3 days. Seedlings were transfer to hydroponic conditions and cultivated for 25 days to the three-leaf one-heart stage; the water was changed every 4 days, and 1/2 Hoagland nutrient solution was used for hydroponics. Seedlings with consistent growth were selected for treatment. For both drought-resistant varieties and drought-sensitive varieties, 17% PEG6000 was used for drought treatment at the three-leaf stage, and the leaves treated at 0 h, 12 h, and 24 h were collected. (Figure 15)Three replicates were performed for each treatment, and the RNAprep Pure polysaccharide polyphenol plant total RNA extraction kit (TIANGEN Co., Ltd., Beijing, China) was used to extract total RNA.

**Figure 15.**
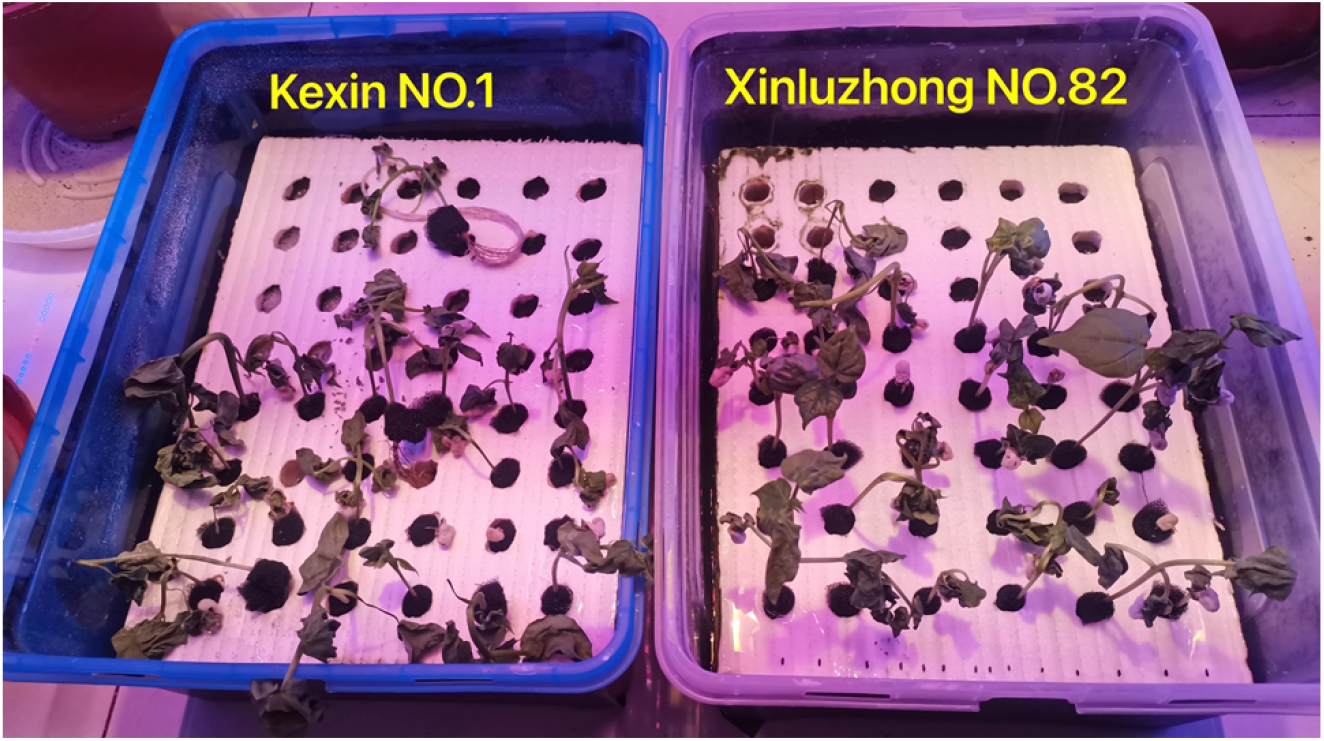
Under seedling drought stress Xinluzhong NO.82 and Kexin NO.1 performance

#### 5.2.2 Cotton transcriptome sequencing

##### (1) Total RNA quality detection

The total RNA quality was determined by UV absorption and denaturing agarose gel electrophoresis.

##### (2) Illumina sequencing

Based on sequencing by synthesis (SBS), the Illumina HiSeq high-throughput sequencing platform sequenced the cDNA library and produced a large amount of high-quality data (raw data).

##### (3) Comparison with reference genome sequences

The genome of *Gossypium hirsutum* was used as a reference for sequence alignment and subsequent analysis. HISAT2 (Li et al., 2014) is an efficient comparison system from RNA sequencing experiment reads, as it had high comparison efficiency.

Download address:UTX_TM1_v2.1.Gossypium_hirsutum.UTX_TM1_v2.1.genome.fa.

##### (4) Bioinformatics analysis

The raw data were filtered, and high-quality clean data were obtained after removing the spliced sequences and low-quality reads; this clean data was used to perform expression analysis and new gene discovery. Advanced analyses, such as the functional annotation and functional enrichment of differentially expressed mRNAs, miRNAs, lncRNAs and circRNAs, were performed according to the expression levels of genes in different samples or groups.

Gene expression has time and space specificity, and external stimuli and the internal environment will affect gene expression. Genes with significantly different expression levels under two different conditions (control and treatment, wild type and mutant type, different time points, different tissues, etc.) are called differentially expressed genes (DEGs). Similarly, transcripts with significantly different expression levels are called differentially expressed transcripts (DETs).

There were two types of differential gene analysis for all samples in this experiment: comparison of differences within groups and comparison of differences between groups. The comparison of intragroup differences refers to the comparison of the same material treated at different times, that is, the comparison of the 16% PEG treatment with the 0 h treatment group at 2 different time points (12 h and 24 h); the comparison of differences between groups refers to the treatment of different materials at the same time, as in the comparisons 0hRvs0hS, 12hRvs12hS, and 24hRvs24hS (Schedule 2).

The gene set obtained by the differential expression analysis is called the differentially expressed gene set and is named using the “A_vs_B” nomenclature. According to the relative level of expression between two (groups) of samples, differentially expressed genes can be divided into upregulated genes and downregulated genes. The expression level of upregulated genes in sample (group) B was higher than that in sample (group) A, while the expression levels of downregulated genes were the opposite. Upregulation and downregulation are relative and are determined by the order of A and B.

##### (5) Analysis of differential gene expression

FPKM (Trapnell C at el., 2010) was used to quantify lncRNA and gene expression as a measure of transcript and gene expression levels. The expression of miRNA was normalized by the TPM (Li et al. 2009) algorithm. The expression levels of circRNAs in each sample were estimated, and junction reads were used to represent the expression levels of circRNAs. Standardized using the SRPBM (Jeck et al.2012) method.

##### (6) Screening of differentially expressed genes

Using DESeq2, discrete convergence evaluation and multiple changes were used to improve the stability and interpretability of the evaluation (Love, 2014). In the detection of differentially expressed genes, fold change (FC) represents the ratio of expression between the two samples (groups). The false discovery rate (FDR) is obtained by correcting the p-value of the difference.

BLAST was used to compare the predicted target gene sequence with the NR (DENG et al. 2006), Swiss-Prot (Apweiler et al. 2004), GO (Michael et al.), COG (Roman et al. 2000), KEGG (Minoru et al. 2004), KOG (Koonin et al. 2004), and Pfam (Eddy. 1998) databases to obtain the annotation information of the target gene.

##### (7) qRT-PCR analysis

Primers were designed by the Primer3 website (Schedule 1), referencing the Tiangen Co., Ltd. (Beijing, China) rapid reverse transcription kit instructions, and amplified using SuperReal PreMix Plus (SYBR Green). qRT-PCR data were analysed by the 2^-ΔΔCt^ method.

## Data availability statements

The raw sequence data reported in this paper have been deposited in NationalCenter for Biotechnology Information(NCBI), under accession number PRJNA769509 and PRJNA769837 that are publicly accessible at https://www.ncbi.nlm.nih.gov/.

## Acknowledge

We would like to thank Xinjiang Academy of Agricultural Sciences, China, for the cotton varieties provided for this study, BMK for the sequencing, and AJE for the English polishing. Thanks to all the people, units and enterprises who have provided help to this study.

## Author Contributions

Zeliang Zhang is the executor of the experimental design and experimental research of this study; Zeliang Zhang, Zhiwei Sang completed the data analysis, the writing of the paper; Junduo Wang, Yajun Liang, Zhaolong Gong participated in experimental design, experimental data collection and test results analysis;Xueyuan Li,Juyun Zheng is the architect and director of the project, guiding experimental design, data analysis, paper writing and modification. The entire authors read and agree to the final text.

## No competing interests among all authors

## Abbreviations

circRNAs: circular RNAs
lncRNAs: long non-coding RNAs
miRNAs: micro RNAs
mRNAs: messenger RNAs
ceRNAs: competing endogenous RNAs
KEGG: kyoto encyclopedia of genes and genomes
DE: differentially expressed
GO: gene ontology
PCA: principal components analysis
GABA: γ-Aminobutyric acid
NAD: nicotinamide adenine dinucleotide
PEG: polyethylene glycol
BP: biological process
CC: cell component
MF: molecular function
qRT-PCR: quantitative Real-Time PCR
TEM: transmission electron microscopy

---

**Schedule 1.**
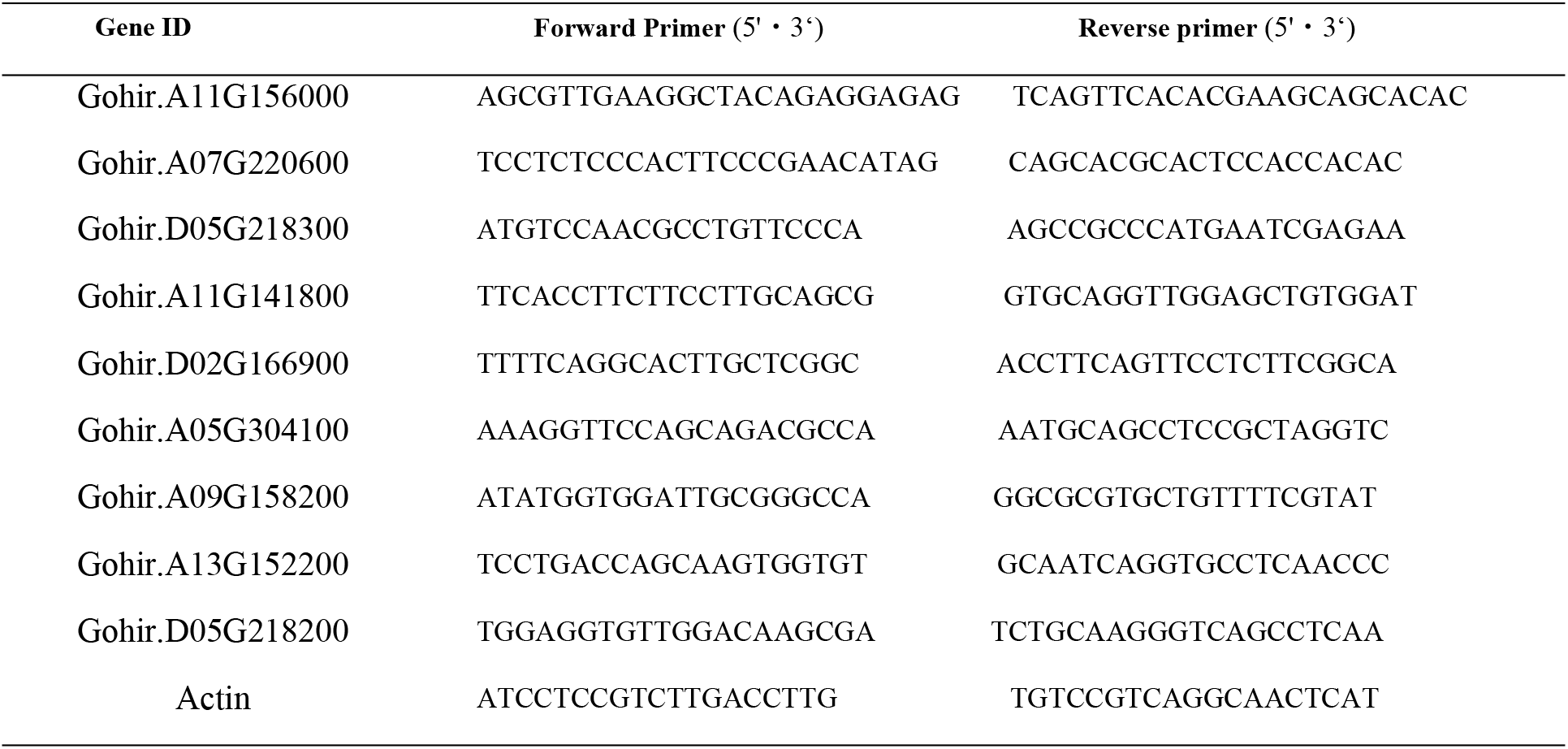
Primer sequences of differentially expressed genes

**Schedule 2.**
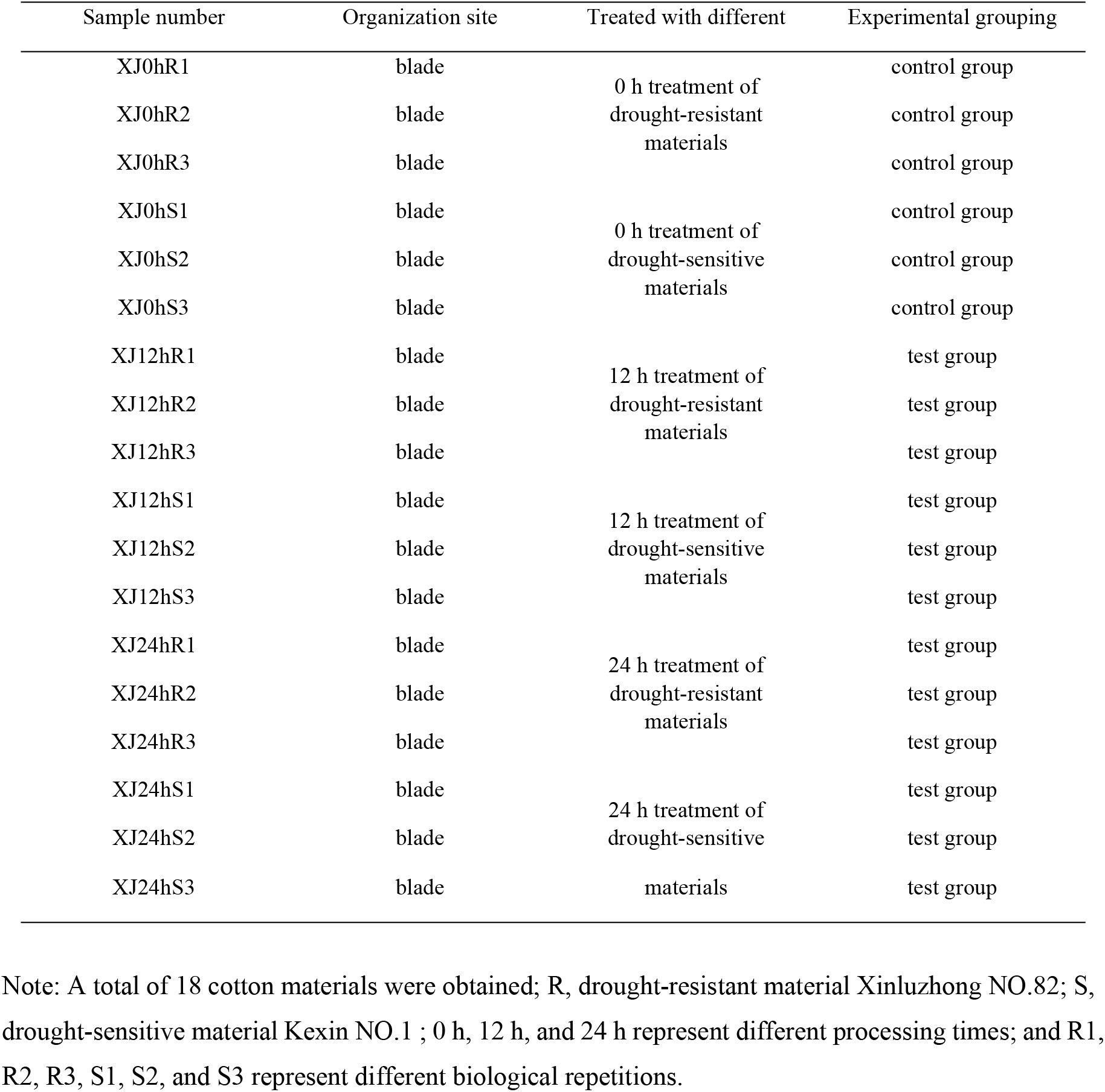
Experimental grouping

**Schedule 3.** Gohir.A11G156000.v2. 1. expression levels

| XJ0h<br>R1_C<br>ount | XJ0h<br>R2_C<br>ount | XJ0h<br>R3_C<br>ount | XJ12<br>hR1_<br>Count | XJ12<br>hR2_<br>Count | XJ12<br>hR3_<br>Count | XJ0h<br>R1_F<br>PKM | XJ0h<br>R2_F<br>PKM | XJ0h<br>R3_F<br>PKM | XJ12h<br>R1_F<br>PKM | XJ12h<br>R2_F<br>PKM | XJ12h<br>R3_F<br>PKM | P-v<br>alu<br>e | lo<br>g2<br>F<br>C | reg<br>ula<br>ted |
| --- | --- | --- | --- | --- | --- | --- | --- | --- | --- | --- | --- | --- | --- | --- |
| 4003 | 2474 | 3860 | 9632 | 7280 | 6983 | 16.35 | 13.45 | 16.82 | 46.05 | 40.35 | 37.97 | 1.2<br>2E<br>-07 | 0.<br>92<br>2 | up |
| XJ0h<br>S1_C<br>ount | XJ0h<br>S2_C<br>ount | XJ0h<br>S3_C<br>ount | XJ12<br>hS1_<br>Count | XJ12<br>hS2_<br>Count | XJ12<br>hS3_<br>Count | XJ0hS<br>1_FP<br>KM | XJ0hS<br>2_FP<br>KM | XJ0hS<br>3_FP<br>KM | XJ12h<br>S1_FP<br>KM | XJ12h<br>S2_FP<br>KM | XJ12h<br>S3_FP<br>KM | P-v<br>alu<br>e | lo<br>g2<br>F<br>C | reg<br>ula<br>ted |
| 3044 | 2876 | 2983 | 6772 | 7408 | 7241 | 16.60 | 14.66 | 15.43 | 35.89 | 40.32 | 38.38 | 1.1<br>2E<br>-12 | 0.<br>78<br>4 | up |
| XJ12<br>hR1_<br>Count | XJ12<br>hR2_<br>Count | XJ12<br>hR3_<br>Count | XJ24<br>hR1_<br>Count | XJ24<br>hR2_<br>Count | XJ24<br>hR3_<br>Count | XJ12h<br>R1_F<br>PKM | XJ12h<br>R2_F<br>PKM | XJ12h<br>R3_F<br>PKM | XJ24h<br>R1_F<br>PKM | XJ24h<br>R2_F<br>PKM | XJ24h<br>R3_F<br>PKM | P-v<br>alu<br>e | lo<br>g2<br>F<br>C | reg<br>ula<br>ted |
| 9632 | 7280 | 6983 | 2018 | 2688 | 3567 | 46.05 | 40.35 | 37.97 | 10.84 | 14.24 | 17.42 | 6.3<br>8E<br>-13 | -1.<br>23<br>9 | do<br>wn |
| XJ12<br>hS1_<br>Count | XJ12<br>hS2_<br>Count | XJ12<br>hS3_<br>Count | XJ24<br>hS1_<br>Count | XJ24<br>hS2_<br>Count | XJ24<br>hS3_<br>Count | XJ12h<br>S1_FP<br>KM | XJ12h<br>S2_FP<br>KM | XJ12h<br>S3_FP<br>KM | XJ24h<br>S1_FP<br>KM | XJ24h<br>S2_FP<br>KM | XJ24h<br>S3_FP<br>KM | P-v<br>alu<br>e | lo<br>g2<br>F<br>C | reg<br>ula<br>ted |
| 6772 | 7408 | 7241 | 2537 | 3302 | 2516 | 35.89 | 40.32 | 38.38 | 13.28 | 17.41 | 12.90 | 1.7<br>4E<br>-12 | -1.<br>10<br>1 | do<br>wn |

**Schedule 4.** Gohir.A07G220600.v2. 1. expression levels

| XJ0h<br>R1_C<br>ount | XJ0h<br>R2_C<br>ount | XJ0h<br>R3_C<br>ount | XJ12<br>hR1_<br>Count | XJ12<br>hR2_<br>Count | XJ12<br>hR3_<br>Count | XJ0h<br>R1_F<br>PKM | XJ0h<br>R2_F<br>PKM | XJ0h<br>R3_F<br>PKM | XJ12h<br>R1_F<br>PKM | XJ12h<br>R2_F<br>PKM | XJ12h<br>R3_F<br>PKM | P-v<br>alu<br>e | log2<br>FC | reg<br>ula<br>ted |
| --- | --- | --- | --- | --- | --- | --- | --- | --- | --- | --- | --- | --- | --- | --- |
| 3279 | 2181 | 3912 | 164 | 287 | 156 | 8.600 | 7.616<br>302 | 10.94<br>6612 | 0.503<br>461 | 1.019<br>193 | 0.542<br>571 | 1.8<br>0E<br>-35 | -4.07<br>8519<br>682 | do<br>wn |
| XJ0h<br>S1_C<br>ount | XJ0h<br>S2_C<br>ount | XJ0h<br>S3_C<br>ount | XJ12<br>hS1_<br>Count | XJ12<br>hS2_<br>Count | XJ12<br>hS3_<br>Count | XJ0h<br>S1_F<br>PKM | XJ0h<br>S2_F<br>PKM | XJ0h<br>S3_F<br>PKM | XJ12h<br>S1_F<br>PKM | XJ12h<br>S2_F<br>PKM | XJ12h<br>S3_F<br>PKM | P-v<br>alu<br>e | log2<br>FC | reg<br>ula<br>ted |
| 2364 | 3325 | 3246 | 370 | 281 | 154 | 8.282 | 10.88<br>4507 | 10.78<br>2553 | 1.259<br>015 | 0.979<br>12 | 0.520<br>93 | 8.7<br>0E<br>-31 | -3.83<br>2399<br>289 | do<br>wn |
| XJ12<br>hR1_<br>Count | XJ12<br>hR2_<br>Count | XJ12<br>hR3_<br>Count | XJ24<br>hR1_<br>Count | XJ24<br>hR2_<br>Count | XJ24<br>hR3_<br>Count | XJ12h<br>R1_F<br>PKM | XJ12h<br>R2_F<br>PKM | XJ12h<br>R3_F<br>PKM | XJ24h<br>R1_F<br>PKM | XJ24h<br>R2_F<br>PKM | XJ24h<br>R3_F<br>PKM | P-v<br>alu<br>e | log2<br>FC | reg<br>ula<br>ted |
| 164 | 287 | 156 | 4256 | 2021 | 2709 | 0.503 | 1.019<br>193 | 0.542<br>571 | 14.68<br>4347 | 6.871<br>942 | 8.496<br>771 | 3.5<br>5E<br>-19 | 4.00<br>3553<br>295 | up |
| XJ12<br>hS1_<br>Count | XJ12<br>hS2_<br>Count | XJ12<br>hS3_<br>Count | XJ24<br>hS1_<br>Count | XJ24<br>hS2_<br>Count | XJ24<br>hS3_<br>Count | XJ12h<br>S1_F<br>PKM | XJ12h<br>S2_F<br>PKM | XJ12h<br>S3_F<br>PKM | XJ24h<br>S1_F<br>PKM | XJ24h<br>S2_F<br>PKM | XJ24h<br>S3_F<br>PKM | P-v<br>alu<br>e | log2<br>FC | reg<br>ula<br>ted |
| 370 | 281 | 154 | 2738 | 3093 | 2151 | 1.259 | 0.979<br>12 | 0.520<br>93 | 9.204<br>302 | 10.47<br>6203 | 7.083<br>376 | 3.8<br>9E<br>-23 | 3.43<br>5904<br>968 | up |

**Attach Figure 1.**
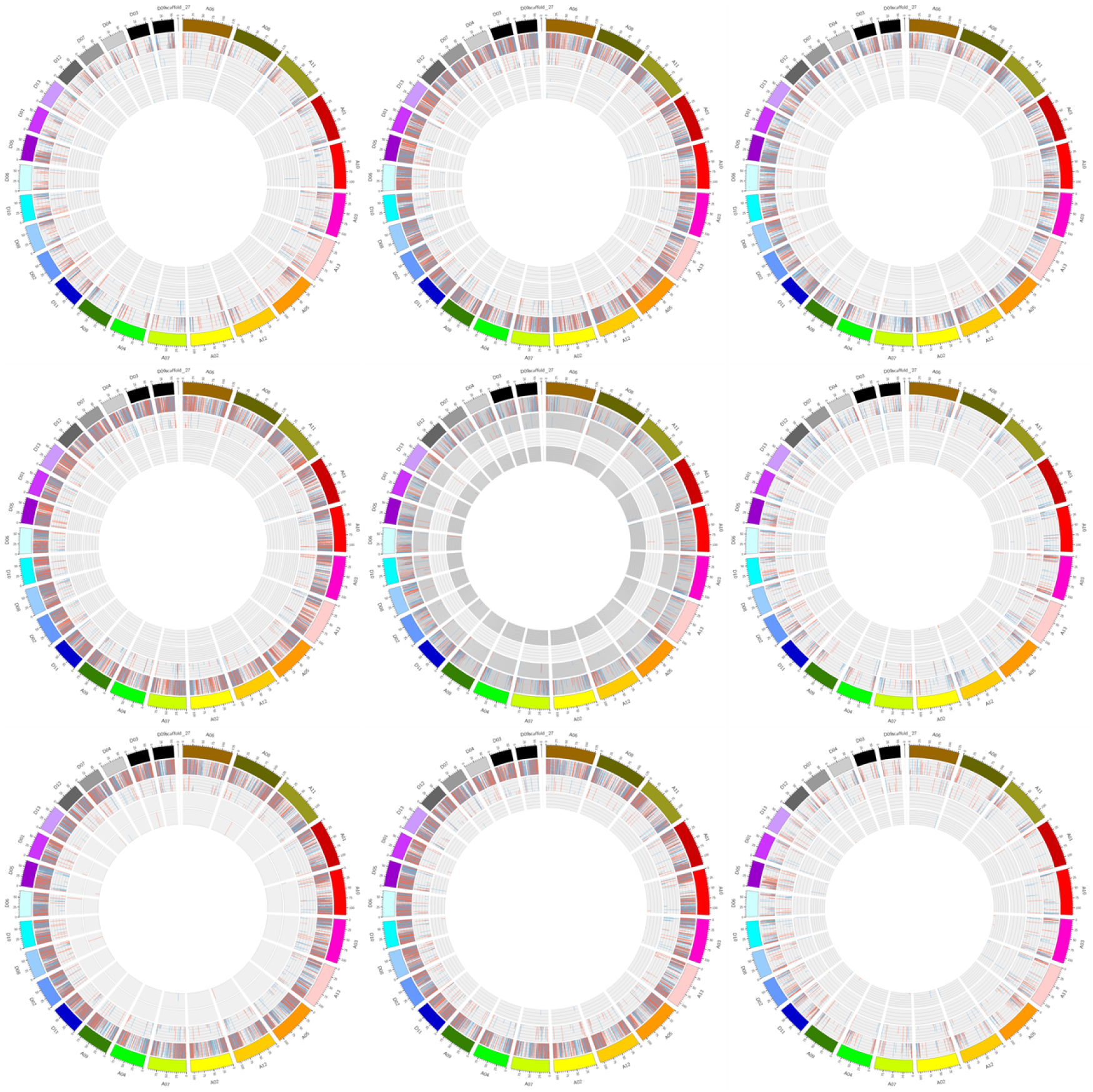
Circos diagrams for different levels of significance across RNA types Note: The outermost circle shows chromosome information, followed by mRNA, lncRNA, circRNA, and miRNA. In the circle diagram of each group, red represents upregulation, blue represents downregulation, and the height represents significance (-log10(FDR or P-value)).

